# Optogenetic Clustering of Human IRE1 Reveals Differential Regulation of Transcription and mRNA Splice Isoform Abundance by the UPR

**DOI:** 10.1101/2025.07.16.665212

**Authors:** Jacob W. Smith, Damien B. Wilburn, Vladislav Belyy

**Affiliations:** The Ohio State Biochemistry Program, The Ohio State University, Columbus, OH 43210; Center for RNA Biology, The Ohio State University, Columbus, OH 43210; Department of Chemistry and Biochemistry, The Ohio State University, Columbus, OH 43210

## Abstract

Inositol-requiring enzyme 1 (IRE1) is one of three known sensor proteins that respond to homeostatic perturbations in the metazoan endoplasmic reticulum. The three sensors collectively initiate an intertwined signaling network called the Unfolded Protein Response (UPR). Although IRE1 plays pivotal roles in human health and development, understanding its specific contributions to the UPR remains a challenge due to signaling crosstalk from the other two stress sensors. To overcome this problem, we engineered a light-activatable version of IRE1 and probed the transcriptomic effects of IRE1 activity in isolation from the other branches of the UPR. We demonstrate that 1) oligomerization alone is sufficient to activate IRE1 in human cells, 2) IRE1’s transcriptional response evolves substantially under prolonged activation, and 3) the UPR induces major changes in mRNA splice isoform abundance in an IRE1-independent manner. Our data reveal previously unknown targets of IRE1 transcriptional regulation and direct degradation. Additionally, the tools developed here will be broadly applicable for precise dissection of signaling networks in diverse cell types, tissues, and organisms.

## Introduction

The endoplasmic reticulum (ER) performs critical roles in eukaryotic cell biology, including protein maturation and lipid metabolism. The dynamic balance between cellular demand and ER capacity is regulated by an intricate signaling network called the Unfolded Protein Response (UPR)^1,2^. The metazoan UPR dynamically integrates the outputs of the three known ER-resident sensor proteins: Protein Kinase R-like ER Kinase (PERK), Activating Transcription Factor 6 (ATF6), and Inositol Requiring Enzyme 1 (IRE1) to direct the cell towards either homeostasis or programmed cell death^3^. The UPR plays a major part both in human development and in diseases such as type 2 diabetes, neurodegeneration, and several types of cancer including multiple myeloma and triple-negative breast cancer^4–10^. Therefore, developing a robust model of UPR signaling is necessary to understand cellular proteostasis, to develop therapeutics against ER-associated diseases^11^, and to learn how to prevent drug-induced ER stress^12,13^. However, building such a model has been complicated by the interconnected nature of the UPR and the experimental difficulties associated with isolating the individual contributions of the three sensor proteins.

IRE1, the most evolutionarily conserved of the three UPR sensors and the focus of this study, is a transmembrane protein composed of a stress-sensing domain within the ER lumen, a single-pass transmembrane domain, and a bifunctional kinase/RNase domain in the cytosol^14–17^. Inactive IRE1α (the primary paralog in mammals^18^; simply IRE1 hereafter) is thought to exist as unphosphorylated monomers or dimers^19^ that further oligomerize in response to ER stress^20,21^. This oligomerization enables *trans*-autophosphorylation of the kinase domain^22^ and subsequent activation of the RNase domain^23^. Fully activated IRE1 recognizes and cleaves the ‘unspliced’ mRNA of the transcription factor X-box binding protein 1 (*XBP1u*) at two locations^24,25^ that are then ligated together by the cytosolic RTCB complex^26,27^ into a form that encodes the ‘spliced’ XBP1 protein (*XBP1s*)^28^. While XBP1u is not thought to be an active transcription factor, XBP1s upregulates hundreds of genes including those encoding critical components of protein-processing machinery in the ER^29–31^. IRE1 has also been reported to cleave other mRNA substrates in a process called Regulated IRE1-Dependent Decay (RIDD)^32,33^ that appears to be impacted by IRE1’s oligomeric and phosphorylation state^34^. RIDD is thought to be a mechanism by which IRE1 can independently augment the slow transcriptional response of XBP1s to rapidly reduce expression of a select group of genes. In addition, IRE1 has been reported to cleave a more extensive pool of RNAs lacking the endomotif characteristic of RIDD targets in a process termed RIDDLE (RIDD lacking endomotif)^34^.

Substantial effort has been invested in understanding IRE1’s specific contributions to the mammalian UPR. Published experimental strategies include ectopic expression of XBP1s^35^, UPR activation with suppression of the IRE1 pathway^36–38^, induction of ER stress with the suppression of the other UPR sensors^39^, and activation of the IRE1/XBP1 axis with small molecules^40–42^. These foundational studies have revealed major conserved effects of IRE1 activation, though every approach comes with caveats: *XBP1u* may not be the only substrate of IRE1^32–34^, so controlling XBP1s expression may not capture the entirety of IRE1 function; acute ER stress may obscure certain effects of IRE1 signaling that occur under the mild levels of ER stress that cells are more likely to encounter *in vivo*; finally, pharmacological activation of IRE1 may induce off-target effects^42^ and does not directly change the oligomeric state of IRE1, which may play a role in its substrate preferences^34^. As a result, despite decades of study, the full effects and regulation of IRE1 signaling are still unclear.

Furthermore, critical aspects of IRE1 regulation remain uninvestigated, such as the temporal dynamics of the IRE1 transcriptional program. The effects of chronic IRE1 activation, as seen in type II diabetes, are easily obscured by laboratory activation of acute ER stress, as IRE1 is known to be attenuated by prolonged PERK activity^43–45^. IRE1 is also involved in functions beyond ER proteostasis, including cell differentiation^46–49^ and regulating lipid imbalances^50^, but the aforementioned experimental challenges have prevented an in-depth characterization of these additional roles. Current research also illustrates the importance of aspects of transcriptional regulation that have been understudied in the context of the UPR, such as transcription termination^51^, alternative splicing^52^, and mRNA stability^53^. To investigate these questions and generate a biologically relevant model for probing strictly IRE1-dependent effects, we sought a way to selectively trigger IRE1 in living cells while mimicking its native mechanism of activation. To this end, we developed a method for controlling the oligomeric transition of IRE1 that enables direct, rapid control over IRE1 activity with limited perturbations to the cellular environment. We then used this approach to reveal previously unappreciated aspects of IRE1’s transcriptional program.

## Results

### Optogenetic Clustering of IRE1 Causes Robust IRE1 Clustering and Activation

To enable minimally invasive, specific activation of IRE1 in live cells, we appended the light-inducible clustering domain CRY2clust^54^ along with the red fluorescent protein mCherry^55^ to the cytosolic C-terminus of the human IRE1 protein lacking the lumenal domain (to prevent activation by ER stress). CRY2clust undergoes rapid, reversible homo-oligomerization in response to illumination with blue light,

Immunoblotting for IRE1 showed that expression levels were unchanged by treatment with light or the ER stress-inducing glycosylation inhibitor tunicamycin (Tm) and that Opto-IRE1 expression was approximately 170 times higher than WT IRE1 levels. Separation of phosphorylation states by Phos-tag PAGE prior to Western blotting indicated that WT IRE1 becomes phosphorylated in response to Tm and not light (Fig. 1B). Inversely, Opto-IRE1 only showed a change in phosphorylation upon light treatment, not Tm. Next, we tested the *XBP1* splicing activity of Opto-IRE1 cells in response to blue LED light and to ER stress induced by treatment with Tm. In sharp contrast to the parental U2-OS Flp-In T-REx cells in which IRE1 had not been knocked out (wild-type, WT, hereafter), the cells expressing Opto-IRE1 exhibited a marked increase in *XBP1* mRNA splicing in response to blue light but not Tm (which was expected since our Opto-IRE1 construct lacks the ER-lumenal sensor domain). The lowest light irradiance to induce maximal splicing was ∼250 µW/cm^2^ (Fig. 1C), and all further experiments were performed at this irradiance to minimize potential off-target effects of illumination. Opto-IRE1 yielded somewhat lower maximal splicing levels than WT IRE1, which we attribute at least partially to the heterogeneous expression levels of Opto-IRE1 that we observed even after clonal selection and FACS. Despite the elevated expression level, baseline *XBP1* splicing by Opto-IRE1 was not above that of WT cells, presumably because the absence of the lumenal domain and its binding interfaces greatly decreases the protein’s constitutive propensity for oligomerization. Live-cell spinning-disk confocal microscopy imaging of these cells revealed that Opto-IRE1 could be clustered by illumination with blue light within minutes to form visibly distinct puncta. These fully disappeared several minutes after blue light was turned off and re-formed in response to subsequent rounds of illumination, demonstrating the speed and reversibility of the process (Fig. 1D). Taken together, our results strongly suggest that oligomerization alone is sufficient in these cells to induce both phosphorylation and RNase activation of IRE1.

**Figure 1.**
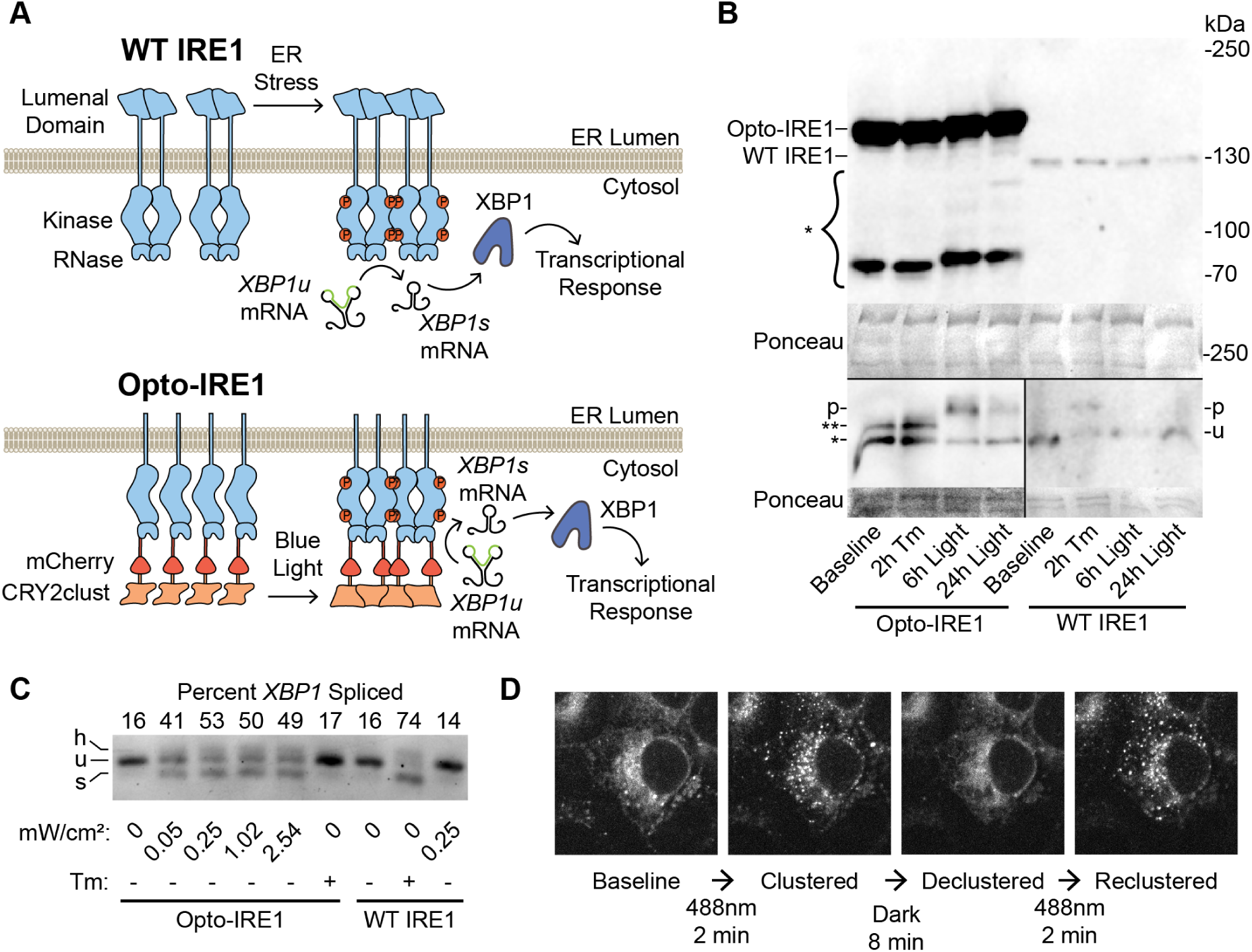
Development and Validation of Opto-IRE1. (A) Diagram of the currently understood mechanism of activation for WT IRE1 (top) and design of our Opto-IRE1 construct (bottom). WT IRE1 and Opto-IRE1 form oligomers in response to ER stress and blue light, respectively, and undergo trans-autophosphorylation and activation to splice *XBP1u* mRNA. (B) Anti-IRE1 Western blots of WT IRE1 and Opto-IRE1 cells treated with tunicamycin (Tm) or light and resolved by SDS-PAGE (top) and Phos-tag SDS-PAGE (bottom). An asterisk denotes several bands for Opto-IRE1 below the full-sized 149kDa band, likely representing truncations of the 58kDa CRY2clust domain. For the Phos-tag gels, the phosphorylated bands are denoted as p, the unphosphorylated WT IRE1 band is denoted as u, and the bands denoted with asterisks are either degradation products or indeterminate phospho-states of Opto-IRE1. Ponceau stained bands are shown as loading controls. (C) Agarose gel depicting *XBP1* mRNA splicing assay of Opto-IRE1 and WT IRE1. Labels below the gel indicate treatments of Tm or varying irradiance of light. Bands corresponding to unspliced *XBP1*, spliced *XBP1*, and their hybrid dimer are denoted with u, s, and h, respectively. (D) Confocal fluorescence microscopy images of Opto-IRE1 cells showing the formation and dissolution of Opto-IRE1 clusters in response to 488nm laser light. and we reasoned that CRY2clust-mediated control of IRE1 oligomerization may serve as an effective “switch” for IRE1’s enzymatic activities (Fig. 1A). Coincidentally, a similar optogenetic IRE1 construct was recently used by Liu et al. to study the coordination of IRE1 to stress granules, though its enzymatic functionality and transcriptional effects were not characterized^56^. We stably integrated our construct, which we call Opto-IRE1, into the tetracycline-inducible safe-harbor locus of U2-OS Flp-In T-REx cells in which expression of endogenous IRE1 had previously been abrogated^57^. Successfully integrated clonal populations were isolated using antibiotic selection followed by fluorescence-activated flow cytometry (FACS) to improve expression homogeneity.

### Light-Induced Clustering of Opto-IRE1 Activates Known IRE1 Target Genes Without Triggering General ER Stress

Having validated our ability to selectively stimulate Opto-IRE1’s kinase and RNase activity with light, we set out to map the specific transcriptional effects of isolated IRE1 activation. To this end, we used long-read Oxford Nanopore Technologies (ONT) sequencing to analyze the transcriptomes of Opto-IRE1 or WT IRE1 cells exposed to light or tunicamycin (Fig. 2A). Briefly, RNA was purified from WT or Opto-IRE1 cells, converted to cDNA by SMARTer-based reverse transcription with barcoded oligo-dT primers for multiplexing, sequencing adapters ligated, and analyzed using an ONT PromethION P2 Solo instrument. The resulting reads were basecalled, demultiplexed, aligned to the human reference genome GRCh38.p14, and analyzed for changes in gene expression and splicing using pyDESeq2^58,59^ (see Methods for details). To verify that this data shows IRE1 activity, we quantified the splicing of *XBP1* in these data by dividing the number of reads for *XBP1* that lacked the intron region by the total reads that spanned the *XBP1* intron (Fig. 2B). This closely matched the PCR analysis in Fig. 1C: WT IRE1 only responded to Tm, and Opto-IRE1 only responded to light. Differential gene expression analysis of WT cells exposed to light also showed no appreciable changes (Fig. 2C), suggesting that 250 µW/cm^2^ is a gentle, noninvasive treatment under our cell culture conditions.

**Figure 2.**
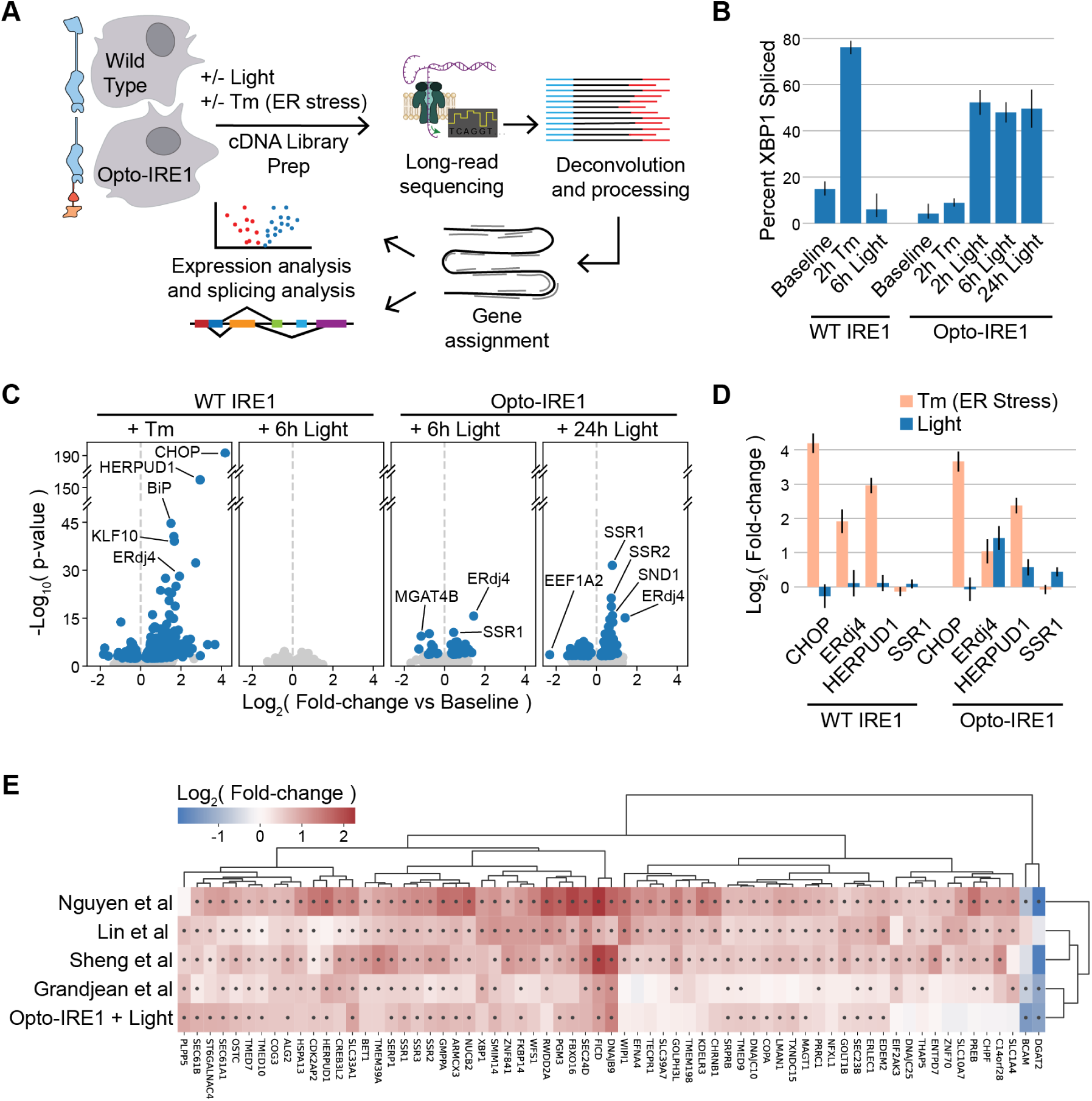
Opto-IRE1 Captures Key Features of the Known IRE1 Transcriptional Program. (A) Schematic depiction of the transcriptomic experiment and analysis. Cells expressing WT IRE1 or Opto-IRE1 are treated with tunicamycin (Tm) or blue light, and their transcriptomes are sampled with long-read nanopore sequencing. This data is then deconvolved, processed, aligned to the GChr38 reference genome, and analyzed to estimate mRNA expression and splicing. (B) Bar plot showing the *XBP1* splicing percentage within the transcriptomic data for WT and Opto-IRE1 cells at baseline and treated with Tm or light. Error bars correspond to 2 standard errors (computed in logit-space) from the mean of three replicates. (C) Volcano plots showing differential gene expression analysis of WT or Opto-IRE1 cells treated with Tm or light. Plots show −log_10_(p-value) (y axis) versus log_2_(fold-change) of gene expression relative to baseline expression (x axis), and genes with FDR-corrected p-value below 0.05 are shown in blue while others are in grey. (D) Bar plot showing log_2_(fold-change of gene expression) from the transcriptomic data of four known UPR targets for WT and Opto-IRE1 cells treated with Tm or light. Error bars are shown corresponding to the mean +/- 2 standard errors from three replicates. (E) Heatmap showing the hierarchically clustered log_2_(fold-change) of genes across four published IRE1 transcriptomic datasets and our dataset for Opto-IRE1 cells with 24 hours light. Black dots indicate FDR-corrected p-values below 0.05.

Our dataset captured the general signature of the UPR and allowed us to validate transcriptional changes that were previously found to be driven predominantly or partially by IRE1 (Fig. 2D): C/EBP homologous protein (CHOP, gene name *DDIT3*), which is known to be affected primarily by PERK activity^60^, was upregulated in both WT and Opto-IRE1 cell lines in response to tunicamycin and not light; ER DNA J domain-containing protein 4 (ERdj4, gene name *DNAJB9*) and Homocysteine inducible ER Protein with Ubiquitin-like Domain 1 (*HERPUD1*), which are known to be upregulated by both IRE1 and ATF6^30,61,62^, showed upregulation in response to both light and tunicamycin in both cell lines; and Signal Sequence Receptor 1 (*SSR1*), which has been shown to be upregulated by IRE1 activity^63,64^, was only upregulated under activation of Opto-IRE1 but not under tunicamycin in either cell line. The observation that SSR1 mRNA levels appear unchanged by tunicamycin treatment of wild-type cells despite IRE1 being demonstrably activated has led us to suspect that the other branches of the UPR may be negating IRE1-mediated regulation of certain genes. Understanding this type of complex co-regulation may become possible with precise and specific activation of IRE1, PERK, and ATF6.

### Opto-IRE1 Elicits a More Limited Transcriptional Program Than Other Methods

To compare our results to other analyses of the transcriptional effects of IRE1 activity, we examined studies with published transcriptomic datasets in human cells that were experimentally comparable with ours. We focused on four studies that examined IRE1’s transcriptional effects by various means: Nguyen et al. infected WT and IRE1-knockout cells with OC43 virus^38^; Sheng et al.^36^ and Lin et al.^37^ used thapsigargin and Tm, respectively, to induce ER stress in WT and *XBP1*-knockdown cells; and Grandjean et al.^41^ treated cells treated with IXA4, a small molecule found to activate IRE1. We reanalyzed the datasets of these studies, selected substantially up- or down-regulated genes, and hierarchically clustered them based on log fold-changes (see Methods for details). Although these data were generated through various methods and in different cell lines, we observed many consistencies between them (Fig. 2E). No genes showed significant contradictory regulation between studies, and some of the most well-established targets of IRE1 (i.e. ERdj4/*DNAJB9*, *FICD*, and *SEC24D*) were consistently upregulated across all five datasets. This demonstrates that IRE1 reliably regulates a core transcriptional program across many biological contexts. The results were almost all upregulated, and only diacylglycerol O-acyltransferase 2 (*DGAT2*) and basal cell adhesion molecule (*BCAM*) were significantly downregulated. While this likely reflects the activating nature of the XBP1s transcription factor, it may also suggest that there is more variation in IRE1-dependent downregulation due to aspects such as cell type. Intriguingly, these are two of the most well-reported RIDD target genes^34^, and they are unregulated or insignificant in the two studies that are XBP1-specific (Sheng et al. and Lin et al.). This finding agrees with XBP1-indendent regulation of these genes. The dataset clustered closest to ours is that of the Grandjean et al. study, which used the small molecule IXA4 to allosterically activate IRE1 trans-autophosphorylation and RNase activity^41,42^. Many genes, such as *EFNA4* and *NFXL1*, showed little to no expression change in our data and the Grandjean et al. data yet showed consistent upregulation in the other three datasets. This may be an indication that some parts of the IRE1 transcriptional program require cooperation of other components of the UPR, since they were only regulated in the datasets that induced ER stress.

### Opto-IRE1 Reveals Temporal Regulation and Novel Transcriptional Targets

Since ER stress *in vivo* varies in duration from transient to chronic^65–68^, we sought to investigate how IRE1’s transcriptional effects evolve over our three timepoints. We found 128 genes that appeared to be regulated (FDR-corrected p-value <= 0.05) by Opto-IRE1 activation under at least one of our timepoints, and we clustered these genes by their log fold-changes (Fig. 3A and Table S1). Nearly all genes were regulated in a consistent manner, with very few showing both up- and down-regulation. Many genes appeared to be delayed until the 6- or 24-hour timepoints, which is potentially a result of chromatin accessibility or regulation by transcription factors downstream from XBP1. Others, such as the GTPase *RALB*, which is thought to mediate ER and Golgi autophagy in connection with cell motility^69,70^, appeared to diminish by 24 hours, potentially representing an early, transient signaling pattern. As expected, the upregulated genes were predominantly involved with various ER functions such as protein folding, modification, glycosylation, and translocation, though the downregulated genes were disproportionately related to transcriptional regulation. Some of the earliest downregulated genes were related to epigenetic identity, such as *KANSL1*^71^ and *MBTD1*^72^. This leads us to suspect that IRE1 activity may dysregulate or alter epigenetic labeling, which could help explain observations that IRE1 activity is necessary for cell differentiation of several cell types^46–49,73^. Broadly, the bias of IRE1 signaling towards upregulation still held true in the temporal analysis, though downregulation of several genes became more prominent at the 24-hour timepoint. Consistent with the inter-study comparison, these downregulated genes showed more variability across time than the upregulated genes. This could be another indication that IRE1-dependent repression is more variable or sensitive to external factors than upregulation.

**Figure 3.**
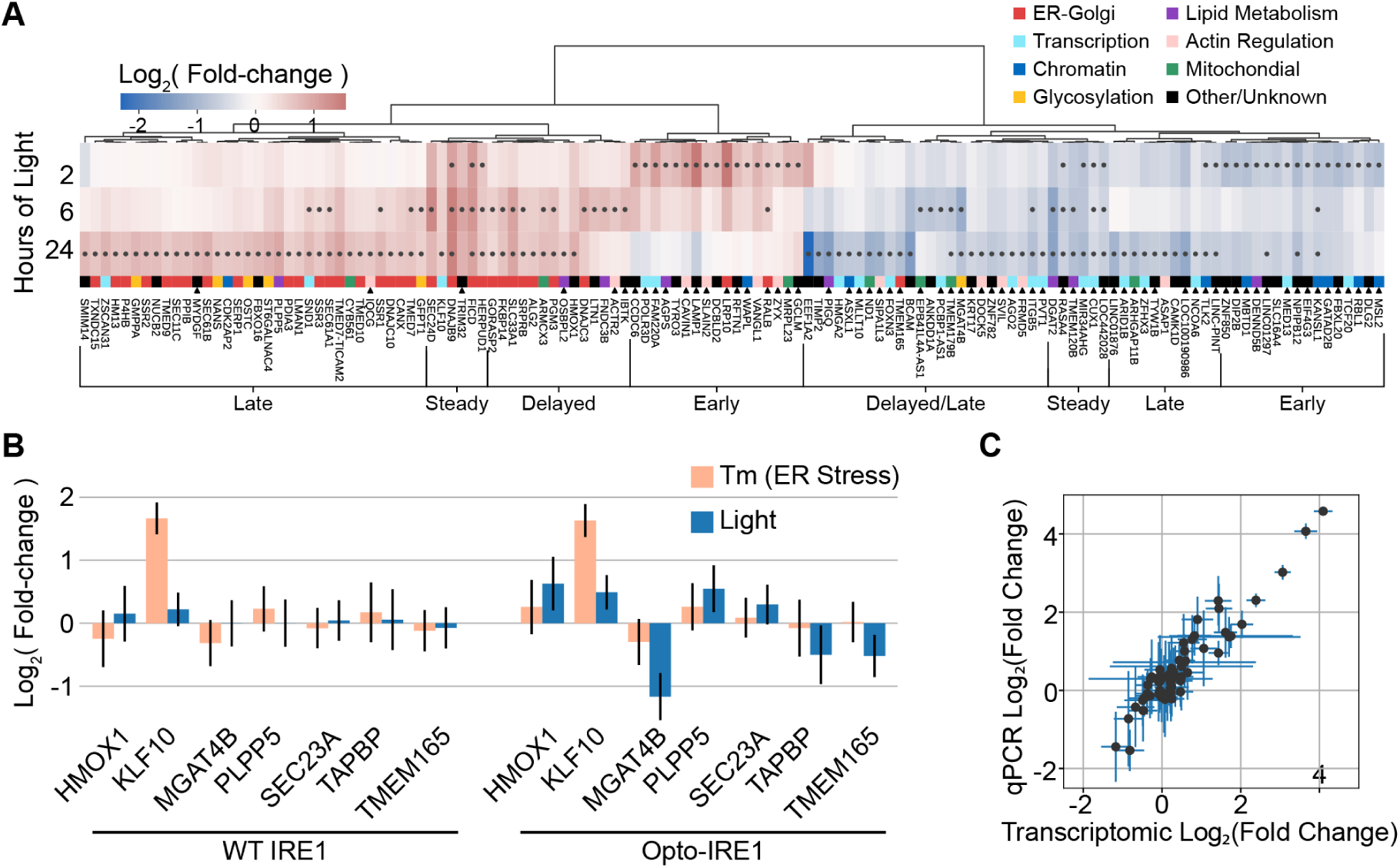
Opto-IRE1 Enables a Nuanced and Temporally Resolved Analysis of IRE1 Signaling. (A) Heatmap showing the hierarchically clustered log_2_(fold-change) of genes across three timepoints of light exposure of Opto-IRE1 cells. Black dots in the heatmap indicate FDR-corrected p-values below 0.05, and the genes are clustered on the log_2_(fold-change) values while the timepoints were kept in chronological order. The larger clusters of genes are annotated based on their temporal expression, and the genes are color-coded based on general cellular function. Black triangles below the function annotations indicate genes that were not found significant in the datasets we compared against. (B) Bar plot showing log_2_(fold-change) of six IRE1 targets for WT cells treated with tunicamycin (Tm) or Opto-IRE1 cells treated with Tm or light. Error bars are shown corresponding to 2 standard errors across three replicates. (C) Scatter plot showing the correlation between log_2_(fold-change) values derived from the transcriptomic data or qPCR assays for 12 genes across 7 experimental conditions. Error bars are given as 2 standard errors across three replicates.

Of the 128 genes in Fig. 3A, 68 did not show changes with FDR-corrected p-value < 0.05 in any of the four transcriptomic datasets discussed above (Table S1). These novel IRE1 target genes are connected to a variety of cellular functions, including glycosylation, mitochondrial function, and transcriptional regulation. The novel downregulated genes disproportionately featured transcription factors, histone modification proteins, and regulators of epigenetic markers, which represents possible mechanisms for how ER stress has been shown to regulate cell identity^67^. Several novel genes were directly connected with ER and Golgi functions: *MGAT4B*, encoding a glycosyltransferase involved in generating branching glycans in the Golgi apparatus^74,75^, and *TMEM165*, encoding a Golgi-resident transmembrane protein thought to regulate calcium and manganese levels in the ER^76,77^, were both downregulated, representing new ways that IRE1 may be regulating glycosylation and ER cation levels, respectively. While it is possible that these genes are artifacts of our specific experimental conditions and cell line, they may have also been identified because inducible oligomerization of IRE1 mimics its endogenous activation in a way that was not fully captured by other approaches.

### The Opto-IRE1 transcriptional program is reproducible and differs from activation by tunicamycin

Several genes appeared to exhibit the same UPR-repressed pattern as *SSR1*, wherein the regulation induced by Opto-IRE1 activation was not observed under activation of the whole UPR with tunicamycin in WT cells: *MGAT4B, TMEM165, HMOX1* (Heme Oxygenase 1, encoding a heme oxygenase that is considered to be anti-inflammatory^78^), *SEC23A* (SEC23 Homolog A, encoding a close paralog of the known IRE1 target *SEC23B*), *PLPP5* (Phospholipid Phosphatase 5, encoding an enzyme involved with phospholipid metabolism^79^), and *TAPBP* (TAP Binding Protein, a component of the protein loading complex^80^). These genes were all regulated almost exclusively by Opto-IRE1, with little to no change in expression during tunicamycin treatment of WT cells (Fig. 3B). In addition to the findings for *SSR1*, these results further illustrate the complexity of the UPR and the nuanced regulation that interferes with isolating the effects of each sensor protein.

To validate some of our results with an orthogonal approach and verify the robustness of our sequencing data analysis, we performed qPCR assays on biological replicates of the experimental conditions used for transcriptomics. For this, we selected the 4 known targets of UPR regulation from Fig. 2C, the 6 UPR-repressed genes discussed above, *XBP1*, and *KLF10* (Kruppel-Like Factor 10, a repressor of TGF-β signaling and another known target of UPR regulation^81^). Overall, our qPCR results agreed remarkably well with our transcriptomic results (Fig. 3C), indicating that Opto-IRE1 induces a reproducible transcriptional program across biological replicates and experimental methods that includes targets that are not detectable upon activation of the broader UPR with tunicamycin.

### The XBP1-independent effects of Opto-IRE1 are limited and directly related to ER function

Many previous studies have observed regulated IRE1-dependent decay (RIDD), which refers to the IRE1-mediated cleavage and subsequent degradation of non-*XBP1* RNAs^32–34,82,83^. To isolate the transcriptomic effects of Opto-IRE1 from those of XBP1, we utilized siRNA to knockdown *XBP1* by approximately tenfold in Opto-IRE1 cells while activating these cells with light (Fig. 4A). We found that the direct effects of Opto-IRE1 activity appeared to be limited relative to the effects of Opto-IRE1 with normal XBP1 expression: only *CD59*, KDEL Receptor 2 (*KDELR2*), Endosulfine Alpha (*ENSA*), and *MGAT4B* appeared to be substantially affected by activation of Opto-IRE1 with an FDR-corrected p-value below 0.05 (Fig. 4B). Given that *ENSA* is upregulated, it is likely not an IRE1 RNase target, which leaves only the three downregulated genes as potential IRE1 substrates. While *CD59* is a commonly reported RIDD target, *MGAT4B* and *KDELR2* have not previously been associated with RIDD and represent novel potential IRE1 RNase substrates. Intriguingly, both genes encode ER-related proteins, suggesting that the effects of RIDD may be partly focused on the ER: as mentioned previously, *MGAT4B* encodes an ER-Golgi glycosyltransferase that is critical for creating branching glycans^74,75^, indicating that RIDD could alter glycosylation patterns by repressing *MGAT4B* (Fig. 4B, C); and *KDELR2* is one of several genes that recognize the KDEL sequence of ER-resident proteins in order to retain them in the ER^84,85^, potentially indicating that RIDD could alter the balance between KDEL-receptors and thus the preference to retain certain KDEL sequences^86^.

**Figure 4.**
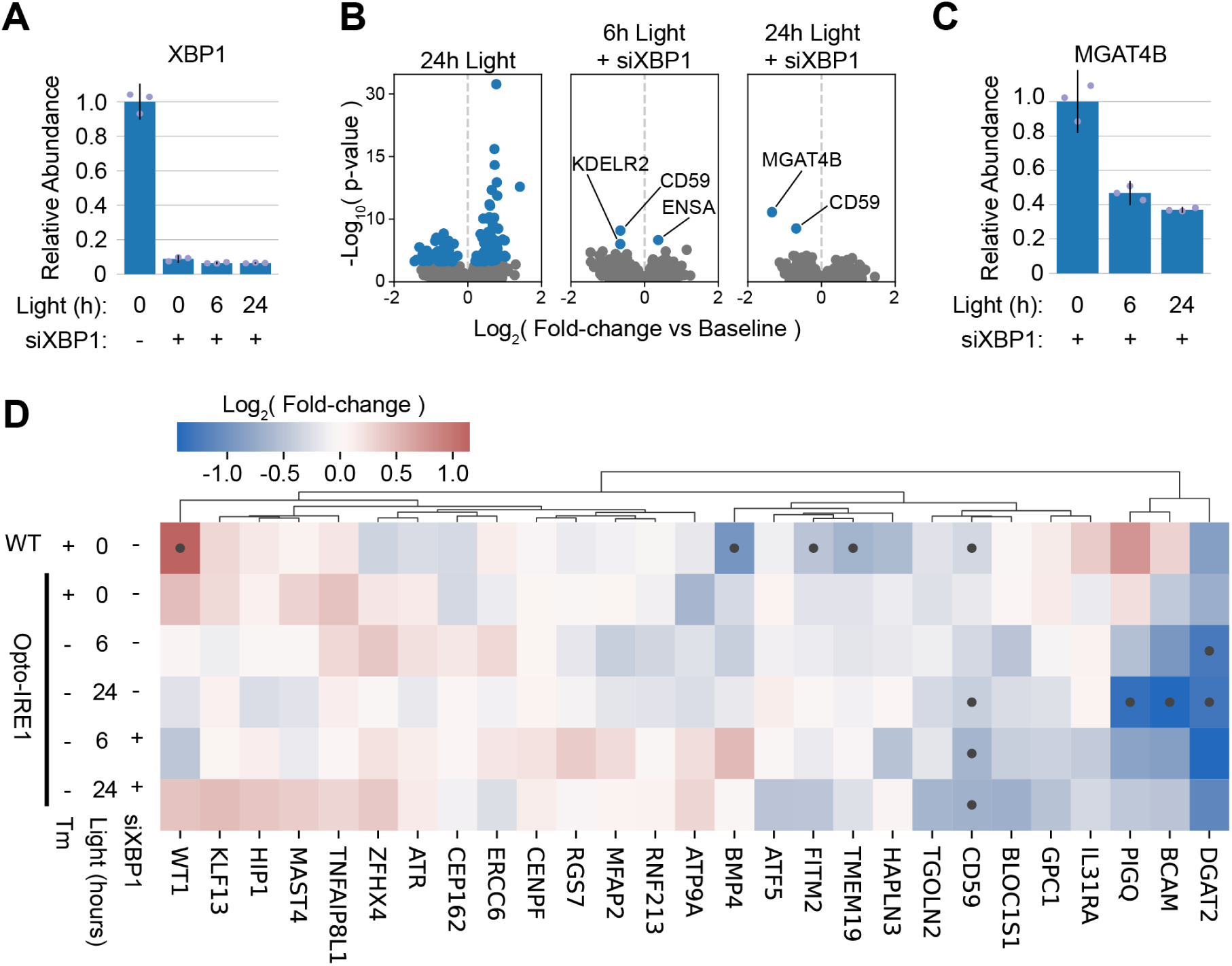
Analysis of XBP1-independent transcriptional effects of Opto-IRE1. (A) Bar plot showing relative qPCR expression levels of *XBP1* in Opto-IRE1 cells treated with light or anti-*XBP1* siRNA (siXBP1). Shown as mean +/- 2 standard errors across three replicates. (B) Volcano plots showing differential gene expression analysis of Opto-IRE1 cells treated with siXBP1 and varying doses of light. Plots show log_10_(p-value) (y axis) versus log_2_(fold-change) of gene expression relative to baseline expression (x axis), and genes with an FDR-corrected p-value below 0.05 are shown in blue while others are in grey. (C) Bar plot showing relative qPCR expression levels of *MGAT4B* in Opto-IRE1 cells treated with light and siXBP1. Shown as averages +/- 2 standard errors across three replicates. (D) Heatmap showing the hierarchically clustered log_2_(fold-change) of previously reported RIDD target genes in WT or Opto-IRE1 cells treated with Tm, light, or siXBP1. Black dots in the heatmap indicate FDR-corrected p-values below 0.05.

We then sought to check whether previously reported RIDD targets were downregulated in our data, so we clustered the log_2_(fold-changes) of genes reported to be XBP1-independent in the Le Thomas et al. and Quwaider et al. studies (Fig. 4D)^34,83^. Few of these genes were notably downregulated under any condition in our data, and only 3 genes were downregulated by 2-fold or more in either of the siXBP1 conditions: *DGAT2, BCAM*, and phosphatidylinositol glycan anchor biosynthesis class Q (*PIGQ)*. Although BCAM^87^ is not strongly associated with ER function, *DGAT2*^88,89^ and PIGQ^90^ are both related to lipid metabolism in the ER. While several other genes (such as *BMP4*, *MFAP2*, and *TMEM19*) were downregulated under Tm treatment of WT cells, their expression did not appear to change under siXBP1 treatment, indicating that these may be regulated by XBP1 rather than directly by RIDD. Altogether, these data suggest that RIDD is a real but limited phenomenon that targets a select set of primarily ER-related gene transcripts for degradation. However, the substrate preference of RIDD has been reported to vary depending on many cellular contexts, including cell type, oligomeric state of IRE1, and PERK interaction^32,34^, so the range of RIDD substrates may well vary by experimental system.

### ER Stress Regulates Abundances of Alternatively Spliced mRNA Isoforms

Since splicing is a critical component of IRE1’s signaling pathway, we wondered whether any other unexpected splicing changes occur under ER stress. After aligning transcripts against the reference genome, we calculated the rate at which each base was retained in the transcriptome as a percent-spliced-in value (%SI) (see Materials and Methods for details) and then compared this %SI value between samples to identify differentially spliced regions. We found that light induced subtle splicing changes in Opto-IRE1 cells, though we did not observe any to the same extent as the *XBP1* intron, supporting the common conclusion that *XBP1* is the only transcript that is directly spliced directly by IRE1 (Fig. 5A). Much more strikingly, tunicamycin-mediated induction of ER stress strongly altered splicing of many regions, including (satisfyingly) the nonconventional intron of *XBP1* (shown in detail in Fig. 5B). We selected two promising alternatively spliced genes to investigate further: Eukaryotic Initiation Factor A2 (*EIF4A2*) encodes one of two isoforms of a key subunit of the eukaryotic initiation factor 4F complex (eIF4F)^91–93^, and Heat Shock Protein 90 B1 (*HSP90B1*) is an ER-resident chaperone that plays key roles in both ER-associated degradation and shuttling of certain proteins through the secretory pathway. For *EIF4A2*, we found that an optional exon was being retained from intron 10 under Tm-induced stress (Fig. 5C), which we verified by PCR and Sanger sequencing across the splice boundary (Fig. 5D, Fig. S4). This exon introduces a premature stop codon that interrupts the C-terminal domain and prevents proper helicase activity, and studies report that the transcript containing this exon is recognized and degraded by the nonsense-mediated decay (NMD) pathway^94,95^. However, the truncated protein appears to be expressed and rapidly ubiquitylated in some cell types^96,97^, and it may play a biologically relevant role^95^. For *HSP90B1*, we found that Tm treatment increased retention of intron 13, which also results in the introduction of a premature stop codon (Fog. 5E, F). The resulting 661aa-long truncated protein, if expressed, would retain the most critical domains for HSP90 chaperone function^98,99^, though it would lack the KDEL ER-retention signal and the C-terminal domain that is thought to regulate co-chaperone interactions^100^. While this isoform may have some function, it is also a candidate for NMD, due to its longer 3’ UTR. The possibility of these transcripts being NMD targets is supported by evidence that PERK activity suppresses the NMD pathway and creates higher levels of NMD-targeted transcripts^101^, which would also result in the accumulation of the alternate *eIF4A2* and HSP90B1 transcripts. Taken together, our analysis supports the existence of uncharacterized aspects of UPR-mediated regulation of protein synthesis that act at the mRNA processing step rather than directly regulating transcription or translation.

**Figure 5.**
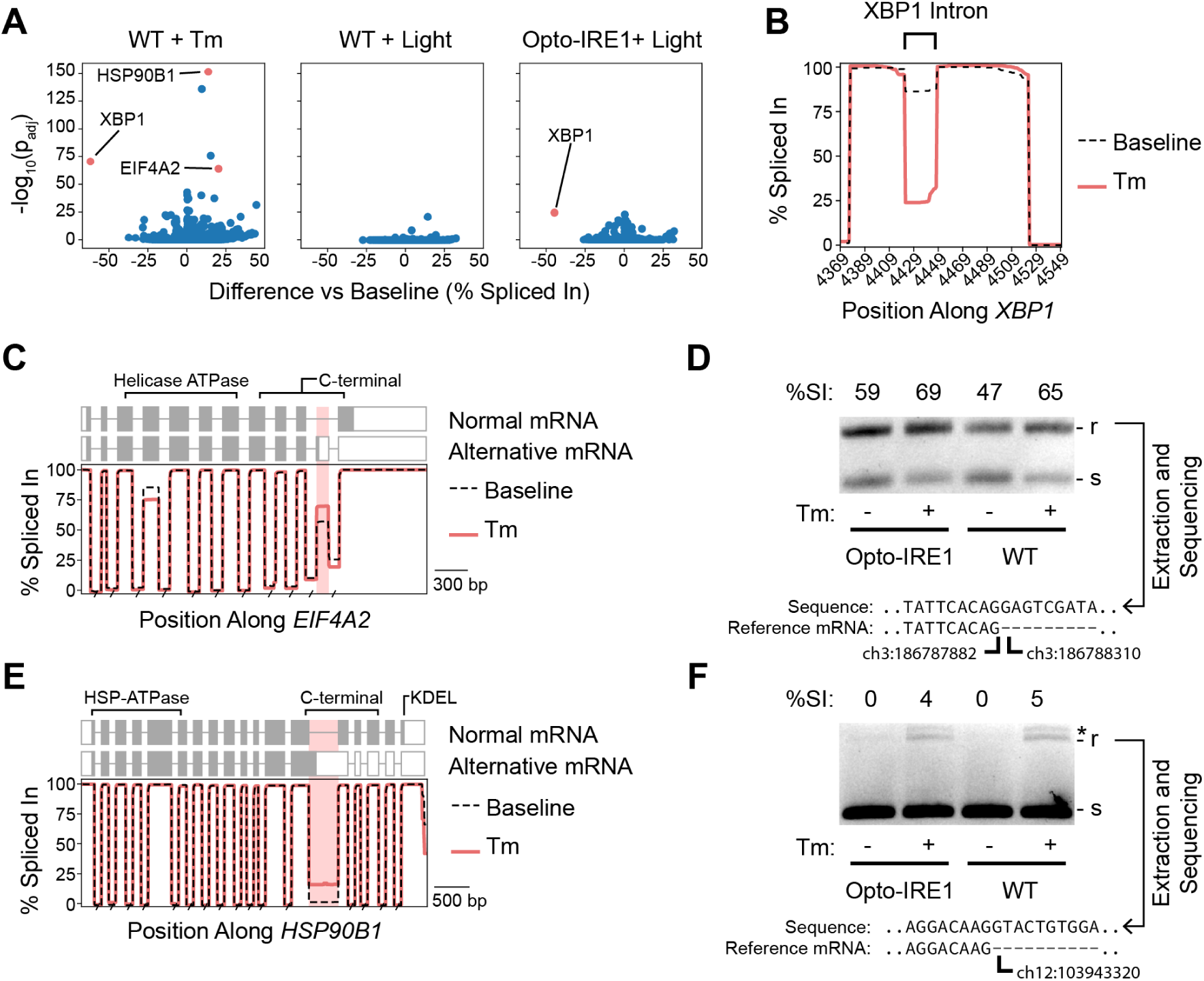
Detection of UPR-dependent alternative splicing. (A) Volcano plots showing the difference in percent-spliced-in of exons relative to baseline (x axis) versus the −log_10_(FDR-corrected p-value) (y axis) for WT cells treated with tunicamycin (Tm) or 6 hours of light and Opto-IRE1 cells treated with 24 hours of light. (B) Line plot showing percent-spliced-in (y axis) versus the genomic position (x axis) along *XBP1* reads near the unconventional intron for WT IRE1 cells at baseline or treated with Tm. (C, E) Line plot showing percent-spliced- in (y axis) versus the genomic position (x axis) along *EIF4A2* reads (C) or *HSP90B1* (E) for WT IRE1 cells at baseline or treated with Tm. The normal and alternative mRNA structures are shown above the plots with boxes indicating exons and the shaded areas indicating the translated reading frame. The known functional protein domains are annotated above these structures. (D, F) Agarose gel depicting the products of a PCR across the alternatively spliced region of *EIF4A2* (D) or *HSP90B1* (F) for WT and Opto-IRE1 cells treated with and without Tm. The larger, intron-retaining product is denoted with ‘r’, the smaller spliced with ‘s’, and off-target products with an asterisk. The sequence of the larger product is shown below, aligned to the 5’ splice junction, and lowercase letters indicate nucleotides that are normally spliced out of the mRNA.

### The UPR Predominantly Alters RNA Isoform Abundances Independently of NMD Inhibition

To gain more insight into the ability of the UPR to alter RNA isoform abundance, we wanted to determine what fraction of our observations in Fig. 5 were due to the known inhibition of NMD by PERK^101^. To do so, we treated WT cells with the SMG1 inhibitor 11j (NMD inhibitor or NMDi)^102^ and analyzed the resulting transcriptome. This revealed that some of the changes in RNA isoform abundance seen under Tm treatment can be replicated by NMDi, though the two treatments are remarkably distinct (Fig. 6A). The NMDi treatment appears to generate more changes in isoform abundance than Tm does, though this method does not discern whether such effects are due to direct NMD inhibition or due to potential off-target effects of the treatment. Intriguingly, the NMD-independent effects of Tm appear to be biased towards retaining RNA regions, whereas the NMDi treatment is relatively unbiased. This may indicate that the UPR selectively alters splicing or splice isoform stability in a manner that favors sequence retention. The change in isoform abundance observed for EIF4A2 appears to be related to NMD, as NMDi replicates and exceeds the splicing pattern seen under Tm treatment (Fig. 6B). However, the change observed for HSP90B1 does not seem to be NMD-dependent, as NMDi did not replicate the increase in sequence retention seen under Tm treatment. This suggests either a direct alteration of the splicing machinery or an altogether different mechanism of RNA splice isoform regulation.

**Figure 6.**
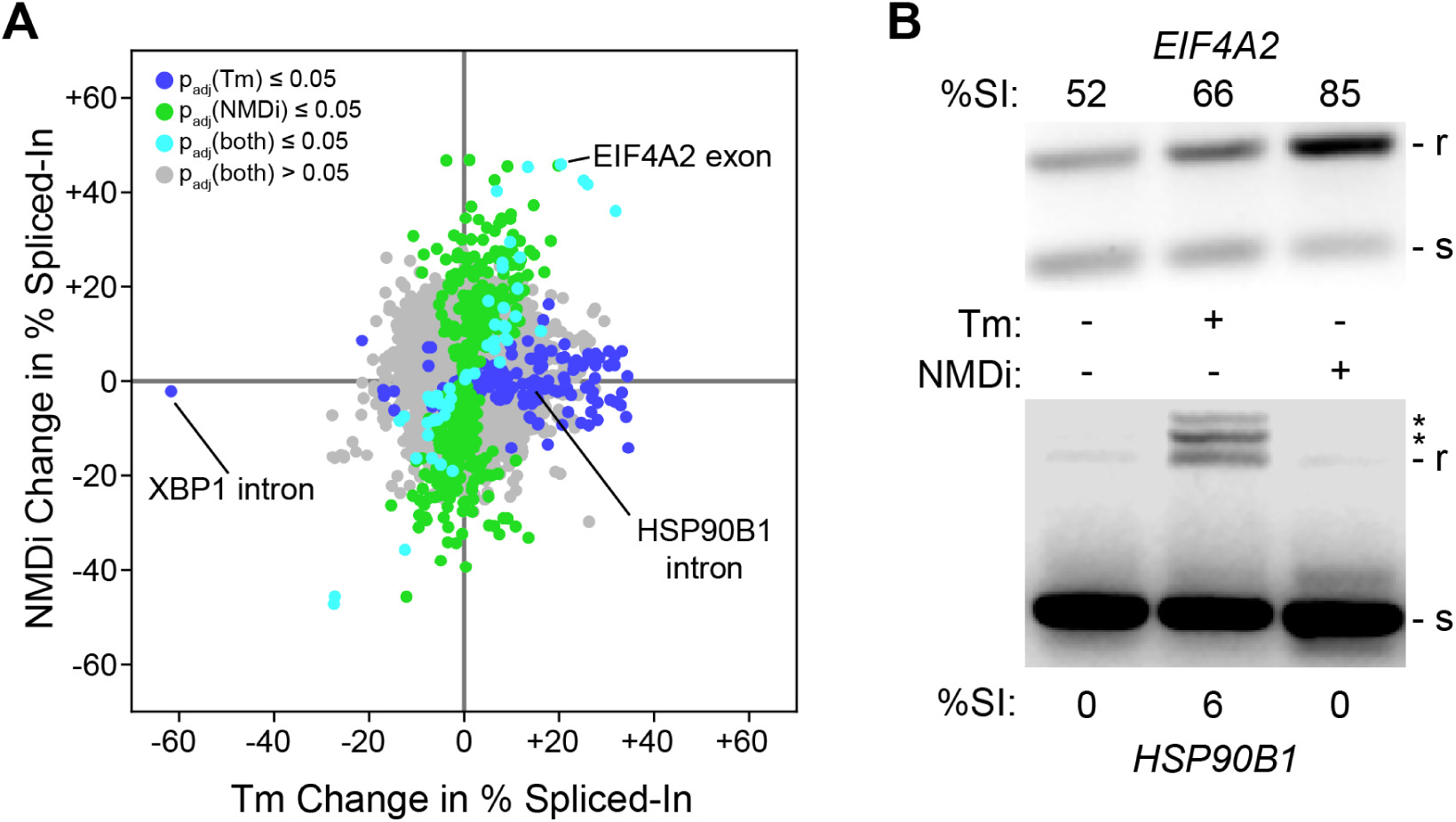
The RNA isoform changes of NMD inhibition overlap with the UPR. (A) A plot of changes in percent spliced in (%SI) paired between treatment with tunicamycin (Tm) and an inhibitor of NMD (NMDi). Each point is an intron/exon region and its color corresponds to the p-value for both treatments. (B) Agarose gel depicting the products of a PCR across the alternatively spliced region of *EIF4A2* (top) or *HSP90B1* (bottom) for WT cells treated with Tm or NMDi. The larger, intron-retaining product is denoted with ‘r’, the smaller spliced with ‘s’, and off-target products with an asterisk.

## Discussion

As a central component of a critical stress response pathway, mammalian IRE1 signaling has been the subject of numerous investigations. While many core components of the IRE1 transcriptional program are generally reproducible and agreed upon, the full extent and temporal progression of IRE1 signaling remain unclear. Major open questions that still evade consensus include the extent of RIDD and the potential interplay between the UPR and mRNA quality control pathways, such as NMD. Moreover, the variation in downstream effects of IRE1 signaling across cell types, tissues, and developmental stages remains poorly understood. Thus, the development of a minimally invasive, easily portable, and highly specific way to induce IRE1 signaling with precise spatiotemporal control promises to greatly enhance our understanding of the UPR’s molecular wiring. The landmark works by Grandjean et al. and Madhavan et al. to develop and characterize the small molecule activator of IRE1, IXA4, made important strides towards filling this gap^41,42^. As demonstrated in this study, Opto-IRE1 exhibits minimal basal activity and does not respond to ER stress due to the absence of its lumenal domain, making it a perfectly orthogonal “switch” that can be integrated into a living system and activated on demand. While small-molecule activation carries obvious pharmacological advantages, our complementary approach in Opto-IRE1 has practically no detectable off-target activity, can be applied with subcellular spatial precision, and is rapidly reversible on the timescale of a few minutes. Since optogenetic control is compatible with multicellular model systems from C. elegans to zebrafish, and mice^103,104^, Opto-IRE1 thus offers a powerful way to study the downstream effects of pulses of IRE1 activation at a precise location and time in the developing or adult organism.

With Opto-IRE1, we have demonstrated that IRE1 signaling exhibits temporal progression with the earliest regulated genes being related to transcription, cell identity, and ER function. This is followed by regulation of many more genes primarily involved in ER function, as well as diverse cellular processes such as lipid metabolism, mitochondrial function, and mRNA regulation. Many of these delayed proteins are likely downstream results of the early elements in the IRE1 pathway, though they may also be inhibited by chromatin accessibility or competing transcription factors. This dynamic regulation is accompanied by early and steady regulation of the most well-known IRE1 targets, such as ERdj4/*DNAJB9* and *HERPUD1*. We have also identified new targets of IRE1 regulation, expanding the known IRE1 transcriptional program. For example, repression of *TMEM165* indicates a potential mechanism for IRE1-mediated control over Ca^2+^ and Mn^2+^ levels in the ER, and repression of *MGAT4B* indicates a new mechanism for modulation of different types of glycosylation. Nearly all of the genes downregulated by Opto-IRE1 in this analysis were not present in the four datasets we compared ours against, which could be a result of several factors. The various experimental methods used between this study and those we re-analyzed could have preferentially captured different gene targets. Alternatively, these genes may not be expressed at measurable levels in the cell lines used by the other studies, thereby preventing observation. Finally, direct activation of IRE1 by inducible oligomerization may capture its endogenous targets in a way that was inaccessible in earlier work.

Beyond direct transcriptional regulation, we also observed ER stress-dependent changes in the abundance of transcript isoforms of several genes, which may result in altered protein levels or functions. These changes can partly be explained as PERK-mediated inhibition of NMD, as they were replicated by treatment with an NMD inhibitor. However, many transcript regions were uniquely regulated by ER stress, potentially representing a new axis by which the UPR regulates protein expression. This could take many forms, such as changes in mRNA quality control in the cytoplasm or altered splicing in the nucleus, and deserves further study.

While the RIDD phenomenon has been heavily investigated, the affected transcripts vary dramatically among studies. Additionally, RIDD has been proposed to be regulated by many factors, including PERK activity and the oligomeric state of IRE1^33,82^. The selectivity and control offered by Opto-IRE1 is very well suited to exploring RIDD and other potential XBP1-independent effects of IRE1 activity. We detected the XBP1-independent downregulation of several previously identified RIDD targets across multiple timepoints, such as *DGAT2*, *BCAM*, and *CD59*^34^ as well as two novel targets *MGAT4B* and *KDELR2* (Fig. 4). Our data suggest that in a scenario where IRE1 alone is activated while other ER stress sensors remain inactive, RIDD may be narrow in scope with very few genes being appreciably repressed.

The new targets of IRE1 regulation that we identified belong to many functional categories, such as transcriptional regulation, lipid metabolism, actin remodeling, and ER homeostasis. Many of the observed downregulated genes, such as KAT8 Regulatory NSL Complex Subunit 1 (*KANSL1*)^71^, were related to epigenetic regulation and chromatin remodeling, indicating that IRE1 may be able to alter epigenetic character. This could be a mechanism for cells to adapt to prolonged ER stress, and may hint at an explanation for IRE1’s control over certain types of cell differentiation^46–49^. We have also further characterized the temporal evolution of IRE1 signaling and showed that the bulk of the transcriptional program is delayed for several hours after IRE1 is activated. This may serve to buffer the cell’s response to low-level ER stress that does not warrant all of IRE1’s transcriptional effects. Consistent with many other studies, our findings place IRE1 at a complex nexus of intracellular signaling, further illustrating the importance of investigating IRE1 activity with precise next-generation approaches.

## Acknowledgements

We thank members of the Belyy and Wilburn laboratories for insights and helpful discussions. We thank Dr. Peter Walter (Altos Labs) and Dr. Diego Acosta-Alvear (Altos Labs) for their thoughtful comments on a draft version of the manuscript. The original plasmid encoding CRY2clust was a kind gift from Won Do Heo (Korea Advanced Institute of Science and Technology, Daejeon, Republic of Korea). This work was supported by NIH R00-GM138896 (to V.B.), NIH R35-GM154813 (to V.B.), NIH R00-HD090201 (to D.B.W.), NIH R35-GM150583 (to D.B.W.), by two Innovation Faculty Startup Awards from JobsOhio and the Enterprise for Innovation, Research and Knowledge (one each to V.B. and D.B.W.) and by a seed grant from the Center for RNA Biology at The Ohio State University (to V.B. and D.B.W.). This work utilized the resources of the Ohio Supercomputer Center^105^ and the OSU Comprehensive Cancer Center shared resources supported by NIH P30CA016058.

## Methods

### Cell culture and experimental reagents

All U2-OS cells were grown at 37°C with 100% humidity and 5% CO_2_ in either DMEM (Gibco) or Fluorobrite DMEM (Gibco), both supplemented with 10% FBS, 2 mM L-glutamine, and 100 units/mL penicillin/streptomycin. Where stated, cells were treated with DMSO-solubilized tunicamycin (Sigma) to 5 ug/mL, DMSO-solubilized doxycycline (Sigma) to 300nM, DMSO-solubilized SMG1 inhibitor 11j (Arctom. CAS NO. 1402452-15-6) to 0.5µM, or blue light treatment with a custom illuminator outputting 250 μW/cm^2^. WT Cells that did not receive tunicamycin of 11j instead received an equivalent dosage of DMSO. All cell lines used in the study tested negative for mycoplasma contamination when assayed with the Universal Mycoplasma Detection Kit (ATCC 301012K) (Fig. S1).

### Generation of the Opto-IRE1 cell line

Opto-IRE1 was expressed in an IRE1-null background using the Flp-In system. The expression plasmid (Addgene plasmid ID 242429) encoding (delta-lumenal domain)IRE1-mCherry-CRY2clust in the pcDNA5/FRT/TO backbone (Thermo Fisher V652020) was stably integrated into IRE1 knock-out U-2 OS Flp-In T-Rex cells that were generated in an earlier study (PWM254)^57^. To achieve stable integration, parental cells were plated into wells of a 6-well plate at a density of 1.7e4 cells/cm^2^. The following day, they were simultaneously transfected with 1.7 μg of the Flp recombinase expression vector, pOG44 (Thermo Fisher V600520), and 300 ng of the IRE1-mCherry-CRY2clust expression plasmid. Transfections were carried out in antibiotic-free DMEM, using the Fugene HD transfection reagent (Promega E2311) and following the manufacturer’s protocol. Twenty-four hours post transfection, cells were split 1:6 into 10-cm dishes and allowed to adhere and recover for an additional 24 hours. The growth medium was then supplemented with 150 μg/mL hygromycin B (Thermo Fisher 10687010) to initiate selection. Cells were maintained in selection medium until single clones became clearly visible and reached ∼3 mm in diameter. Three clones were picked using sterile cloning cylinders (Corning 3166-10), expanded, and assayed for doxycycline-inducible expression of the recombinant construct. The clone that showed the most robust increase in IRE1-mCherry-CRY2 expression upon addition of doxycycline was expanded further and sorted on a Becton Dickinson FACSAria III sorter, selecting for cells with ‘low’ or ‘high’ levels of mCherry fluorescence relative to the parental PWM254 cell line. The ‘low’ pool of sorted cell cells was assigned the identifier “cVBL001”, confirmed to be free of mycoplasma contamination by PCR (30-1012K, ATCC) and used as the “Opto-IRE1” cells for subsequent experiments in this study.

### RNA purification

For all experiments using RNA, cells were seeded into a black 24-well glass-bottom plate at 20,000 cells per well and incubated for 24 hours in supplemented DMEM, then for 24 hours in supplemented Fluorobrite medium with doxycycline. Cells were then given treatments of light, tunicamycin, or SMG1 inhibitor 11j for prescribed durations and irradiances before being washed with PBS and lysed with Trizol (Thermo Fisher). Chloroform was added and RNA was then purified from the aqueous layer using a spin-column kit (RNA Clean & Concentrator-5, Zymo Research #R1015) according to manufacturer protocol and quantified using a Nanodrop One C (Thermo Scientific).

### Anti-XBP1 siRNA treatment

Opto-IRE1 cells were electroporated with 5 µM anti-XBP1 siRNA (Santa Cruz Biotechnology) with a Neon NxT electroporator (Thermo Fisher) set to 1230V, 10ms, and 4 pulses. These cells were then plated for RNA experiments as described above, but with 40,000 cells per well to account for roughly 50% mortality after electroporation.

### Transcriptomic prep

RNA was collected as described above in biological triplicate and converted to double stranded complementary DNA (cDNA) using an adaptation of SMARTer-based protocol^106^. 200 ng total RNA in 11.5 μl water was mixed with 1 μl of 10 μM barcoded poly-dT primer and 1 μl of 10 mM dNTPs before being incubated at 65°C for 5 min, 4°C for 5 min, and 42°C for 2 min. The reaction was then quickly mixed with 0.5 µl RNaseOUT (Thermo Fisher #10777019), 1 µl of 20mM strand-switching primer, 1 µl Maxima H Minus Reverse Transcriptase (Thermo Fisher EP0751), and 4 µl Maxima H Minus buffer. This reaction was incubated at 42°C for 90 min before being heat killed at 85°C for 5 min and cooled to 4°C. This was PCR amplified by combining 2 µl of cDNA, 2 µl Platinum SuperFi II DNA Polymerase (Invitrogen # 12361010), 20 uL Platinum SuperFi II buffer, 2 μl of 12 µM PR2 primer, 2 μl 10 mM dNTPs, and 72 μl of water, and then thermally cycled using the following program: 98°C for 30s, 16x[98°C for 10s, 60°C for 10s, 72°C for 6 min], and held at 4°C. An equal volume of sparQ PureMag Beads was then mixed with the PCR reaction containing amplified cDNA, mixed for 5 min, separated from supernatant on a magnet, washed twice with 500 μl of 70% ethanol, lightly dried, resuspended in 21 μl water, mixed for 10 min, and separated with a magnet. The supernatant containing double stranded cDNA was then quantified by fluorescence using the Quant-iT Picogreen reagent (Thermo Fisher P7589), and equal masses of DNA were pooled from samples with unique molecular barcodes to a total of 150 ng of multiplexed DNA and diluted to 50 μl. 3 μl of enzyme and 7 μl of buffer from the NEBNext Ultra II End Repair/dA-Tailing Module (New England Biolabs E7546S) were added to the DNA pool and incubated at 20°C for 5 min and 65°C for 5 min. End-repaired DNA was then mixed with60μl sparQ PureMag Beads, mixed for 5 min, separated from supernatant on a magnet, washed twice with 200 μl 70% ethanol, lightly dried, resuspended in 60 μl water, mixed for 10 min, and separated with a magnet. The end-repaired cDNA in the supernatant was then ligated to sequencing adapters by adding 25 μl Ligation Buffer and 5 μl Ligation Adapter from Ligation Sequencing Kit V14 (Oxford Nanopore Technologies SQK-LSK114) as well as 10 μl Quick T4 Ligase (New England Biolabs E6056S) before incubation for 10 min at room temperature. 40 μl sparQ PureMag Beads was then added to the ligated DNA library, mixed for 5 min, separated from supernatant on a magnet, washed twice with 250 μl of Short Fragment Buffer, lightly dried, resuspended in 32 μl Elution Buffer (Oxford Nanopore Technologies SQK-LSK114), mixed for 10 min, and separated with a magnet. The resulting supernatant containing the prepared DNA library was then loaded onto PromethION Flow Cell R10.4.1 (Oxford Nanopore Technologies FLO-PRO114M) as per manufacturer protocol and sequenced for 72 hours.

### Transcriptomic Data Analysis

Sequencing traces were basecalled using Dorado v0.9.6 with the dna_r10.4.1_e8.2_400bps_sup v5.0.0 model. The resulting pooled reads were assigned to samples by detecting the barcode sequences of the VN primers: This was done by first searching for the poly-T region of the VN primer and the common primer (CP) sequence, then searching near those features for the unique sequences. Barcodes were confidently assigned when a unique sequence was matched with 3 or fewer mismatches in the correct direction relative to the poly-T region and/or the common sequence, and reads were excluded from further analysis if multiple or no barcodes were successfully assigned. The edlib package was used for all barcode alignments^107^. Any identified barcode, poly-T, and CP sequences were trimmed, and the remaining internal sequence was aligned to the human reference genome GRCh38.p14 (Genome Research Consortium), excluding revised scaffold contigs, using the minimap2 package set to the ‘splice’ preset^108^. The resulting alignments were each assigned to a gene by finding the annotation in the GRCh38 gene file with the greatest ratio of overlapping sequence to the length of the annotation or alignment, whichever is longer. These gene assignments were then used to generate per-sample gene counts, and these counts were analyzed for differential gene expression.

We analyzed the data of our study and those we compared against using the pyDESeq implementation of DESeq2^58,59^. For these analyses, each combination of cell type and treatment was treated as a separate experimental condition, the min_mu parameter was set to 15, and all other parameters were kept as the default values. Genes were selected as ‘significant’ if they had FDR-corrected p-values (Benjamini-Hochberg) ≤0.05 and |log_2_(Fold-Change)| ≥ 0.5.

### Transcriptomic Splicing Analysis

Reads were grouped by gene assignment for each sample, and the CIGAR sequences for their alignments were then used to generate a measure of splicing, wherein each position was defined by the number of spliced-in reads that included a base at the position (including mismatched bases) and the total number of reads that spanned across the position. Short regions in these CIGAR sequences (less than 10 bases) were removed by combining them with the preceding region, and insertions were ignored. To exclude a subset of reads that were found to be short, spurious alignments, reads that were more than 80% aligned to the reference were excluded. The first and last regions of each read were also ignored, which reduced the noise introduced by the variability in alignment of the 3’ and 5’ ends of reads. For each set of biological replicates, these splicing pileups were combined and grouped into distinct regions by calculating the cumulative average spliced-in fraction along the genomic index and starting a new region whenever the fraction differed from the cumulative average by more than 0.1. Additionally, genomic positions were excluded when the total read coverage fell below 5 for any individual sample or below 20 for the average coverage among biological replicates. These regions were then filtered to exclude regions that were not retained in any condition (regions with a mean splicing fraction less than 0.05 for all samples being analyzed). For each set of biological replicates, the number for total reads and spliced-in reads were both averaged across each region and rounded to integers. To estimate the retention rate for each region per experimental condition, a maximum likelihood estimate was calculated for the probability of being retained by minimizing the negative log likelihood function of the data on a negative binomial distribution. For each comparison between conditions, a maximum likelihood estimate was also calculated for the retention rate of the region for all samples between both conditions. To estimate per-region significances for each comparison, the probability of both samples having the same splicing rate was calculated by dividing the likelihood function of the negative binomial distribution fit to the data of both conditions by the product of the likelihood functions of the distributions fit to each condition separately.

### XBP1 splicing assay

RNA was collected as described above and then reverse-transcribed using SuperScript VILO Master Mix (Thermo Fisher #11755050). The resulting cDNA was diluted 1:10 and amplified by PCR with Taq polymerase (Thermo Fisher #10342020) using primers for the XBP1 intron region (see table S1) using the following thermocycler program: 95°C for 1 min, 34x[ 95°C for 30s, 58°C for 30s, 72°C for 30s]. PCR products were separated on a 3% agarose gel stained with SYBR Safe (Thermo Fisher S33102) and imaged on an Azure Biosystems 300 imager. The intensities of the bands for XBP1u, XBP1s, and their hybrid dimer product were each quantified and corrected against the local background gel intensity using ImageJ, and XBP1 splicing was calculated as (XBP1s + hybrid/2) / (XBP1u + XBP1s + hybrid).

### qPCR

RNA was collected as described above in biological triplicate and then reverse-transcribed using SuperScript VILO Master Mix (Thermo Fisher #11755050). The resulting cDNA was diluted 1:10 and used as template for qPCR using PowerUp SYBR Green Master Mix (Thermo Fisher A25741) and 500nM primers (see table S1). This mix was then run in duplicate on either an Applied Biosystems 7500 Fast RT-qPCR or an Azure Biosystems Cielo RT-qPCR with the following qPCR program: 95°C for 5 min, 40x[ 95°C for 30s, 58°C for 30s, 72°C for 30s, measure fluorescence]. The resulting fluorescence data was then used to fit a mathematical model of PCR amplification^109^ in a custom python script to estimate the starting quantity of template in arbitrary units. This estimate was then averaged between the technical duplicates, normalized against the averaged estimate of beta-Actin from the same cDNA, and divided by the Actin-normalized value for the baseline sample. Lastly, the mean and sample standard error of this ratio was calculated across the biological replicates.

### Splicing PCR

RNA was collected as described above and then reverse-transcribed using SuperScript VILO Master Mix (Thermo Fisher #11755050). The resulting cDNA was diluted 1:10 and amplified by PCR using Phusion polymerase (New England Biolabs M0530S) with 500nM primers flanking the alternatively spliced region of each gene (see table XX) using the following thermocycler program: 98°C for 1 min, 35x[ 98°C for 10s, 56°C for 30s, 72°C for 15s], 72°C for 2 min. PCR products were separated on a 3% agarose gel stained with SYBR Safe (Thermo Fisher S33102) and imaged on an Azure Biosystems 300 imager. The intensities of the bands for the spliced and unspliced products were each quantified and corrected against the local background gel intensity using ImageJ.

### Western Blots

Cells were plated at 1.5e5 cells per sample and incubated for 24 hours in supplemented DMEM, then for 24 hours in supplemented Fluorobrite medium with 300 nM doxycycline, then given treatments of light or tunicamycin before being washed and scraped into cold PBS. These cells were then centrifuged and resuspended in 60 μl lysis buffer (50mM Tris, 150mM NaCl, 0.15% SDS, 1% Triton-X, 1mM PMSF, and PhosStop phosphatase inhibitor (Roche)), incubated for 45 minutes on ice, and centrifuged at 2e4 RCF for 5 minutes. Protein in the supernatant was quantified by Pierce BCA assay (Thermo Fisher #A65453), diluted to 1 μg/μl, and 12 μl was boiled for 5 minutes with 3 μL loading dye (30% glycerol, 200mM Tris pH 8.0, 10% SDS, 0.01% bromophenol blue, 40mM DTT). For the normal PAGE, samples were loaded into 5% SDS-PAGE gel, separated by electrophoresis (120V, 120 min, 4°C), and transferred to a nitrocellulose membrane via wet blotting (120V, 90 min, 4°C). For the Phos-tag, lysate was loaded into 5% SDS-PAGE gel with 25μM Phos-tag (NARD Institute AAL-107) and 50μM MnCl_2_, separated by electrophoresis (120V, 140 min, 4°C), and transferred to a nitrocellulose membrane via wet blotting (120V, 150 min, 4°C). The membranes were blocked with 5% fat-free milk powder in TBST (20 mM Tris, 150 mM NaCl, 0.1% Tween 20, pH 7.6) for 1h at room temperature, then incubated with primary antibodies in 5% milk/TBST overnight at 4 °C with agitation. The membranes were washed 3x with TBST, incubated with the secondary antibodies in 5% milk/TBST for 1h at room temperature, washed 3x with TBST, developed with SuperSignal West Femto (Thermo Fisher #34095), and imaged on an Azure Biosystems 300 imager for 10 minutes.

## Data Availability

The code used to process and analyze the raw transcriptomic and qPCR data and to generate the figures in this paper is freely available through Zenodo (https://doi.org/10.5281/zenodo.15988459), along with the source qPCR data, raw gel and blot images, and raw microscopy data. Instructions for how to use the python scripts are in the included README.md file and detailed comments throughout the scripts and IPython notebooks. Design files for the cell illuminator are also available through Zenodo, in the same repostory. De-multiplexed transcriptomic data and per-sample gene counts are deposited in the public National Center for Biotechnology Information Gene Expression Omnibus repository under the data identifier GSE302406. Raw nanopore data and the cell lines used in this paper are available upon request.

