## Supplementary Tables and Figures for "Optogenetic Clustering of Human IRE1 Reveals Differential Regulation of Transcription and mRNA Splice Isoform Abundance by the UPR"

**Supplemental Materials**

| Primer ID | Sequence |
|---|---|
| dT_BC1001_PB | AAGCAGTGGTATCAACGCAGAGTACCACATATCAGAGTGCGTTTTTTTTT TTTTTTTTTTTTTTTTTTTTTVN |
| dT_BC1002_PB | AAGCAGTGGTATCAACGCAGAGTACACACACAGACTGTGAGTTTTTTTT TTTTTTTTTTTTTTTTTTTTTTVN |
| dT_BC1003_PB | AAGCAGTGGTATCAACGCAGAGTACACACATCTCGTGAGAGTTTTTTTTT TTTTTTTTTTTTTTTTTTTTTVN |
| dT_BC1004_PB | AAGCAGTGGTATCAACGCAGAGTACCACGCACACACGCGCGTTTTTTTT TTTTTTTTTTTTTTTTTTTTTTVN |
| dT_BC1005_PB | AAGCAGTGGTATCAACGCAGAGTACCACTCGACTCTCGCGTTTTTTTTTT TTTTTTTTTTTTTTTTTTTTTVN |
| dT_BC1006_PB | AAGCAGTGGTATCAACGCAGAGTACCATATATATCAGCTGTTTTTTTTTTT TTTTTTTTTTTTTTTTTTTTVN |
| dT_BC1007_PB | AAGCAGTGGTATCAACGCAGAGTACTCTGTATCTCTATGTGTTTTTTTTTT TTTTTTTTTTTTTTTTTTTTVN |
| dT_BC1008_PB | AAGCAGTGGTATCAACGCAGAGTACACAGTCGAGCGCTGCGTTTTTTTT TTTTTTTTTTTTTTTTTTTTTTVN |
| dT_BC1009_PB | AAGCAGTGGTATCAACGCAGAGTACACACACGCGAGACAGATTTTTTTT TTTTTTTTTTTTTTTTTTTTTTVN |
| dT_BC1010_PB | AAGCAGTGGTATCAACGCAGAGTACACGCGCTATCTCAGAGTTTTTTTTT TTTTTTTTTTTTTTTTTTTTTVN |
| dT_BC1011_PB | AAGCAGTGGTATCAACGCAGAGTACCTATACGTATATCTATTTTTTTTTTTT TTTTTTTTTTTTTTTTTTTVN |
| dT_BC1012_PB | AAGCAGTGGTATCAACGCAGAGTACACACTAGATCGCGTGTTTTTTTTTT TTTTTTTTTTTTTTTTTTTTTVN |
| strand-switching primer | AAGCAGTGGTATCAACGCAGAGTAC-r(GGG) |
| PR2 primer | AAGCAGTGGTATCAACGCAGAGTAC |
| oJS023_HERPUD1_mRNA_f | TGAGCAGATTCCTCATGGTC |
| oJS024_HERPUD1_mRNA_r | GATCAGTGCCTTCCTGTAAGT |
| oJS027_SSR1_mRNA_f | TGAACCCACAGATTTGGTAGAA |
| oJS028_SSR1_mRNA_r | TGTTGGTAAAGCCTACCAGG |
| oJS031_SEC23A_mRNA_f | TTGACACTGAACATGGAGGC |
| oJS032_SEC23A_mRNA_r | TGCTCCAGACTCCTGCC |
| oJS039_TAPBP_mRNA_f | TATCTCAGTGTACACGACCCC |
| oJS040_TAPBP_mRNA_r | GGCTCATCTCGCAGTGTG |
| oJS051_KLF10_mRNA_f | GTCGAGTGTCTCCGTGC |
| oJS052_KLF10_mRNA_r | ATTTCCATTCTTTCCTCCGCA |
| oJS095_EIF4A2_exon_10_f | CAACAAGTGTCTTTGGTTAT |
| oJS096_EIF4A2_exon_11_r | AAATCGACCCCCTCT |
| oJS099_HSP90B1_exon_13_f | GAAAACTAAGGAGAGTCGT |
| oJS100_HSP90B1_exon_14_r | CTTTCATGATTCTCTCCAT |

| | |
|---|---|
| oJS107_HMOX1_mRNA_f1 | GTGCCACCAAGTTCAAGCAG |
| oJS108_HMOX1_mRNA_r1 | GCAACTCCTCAAAGAGCTGGA |
| oJS123_PLPP5_mRNA_f1 | CGGCCTTCCTGGTGACG |
| oJS124_PLPP5_mRNA_r1 | CGGCTTGGTGGGGAAATACT |
| oJS131_TMEM165_mRNA_f1 | GGCCCGGGTCGAGAAAAT |
| oJS132_TMEM165_mRNA_r1 | TCAGGCGGTTATAGCGCATT |
| oJS165_MGAT4B_mRNA_f4 | CTGCTGCTCTTCTGCCTGT |
| oJS166_MGAT4B_mRNA_r4 | CTTTCTGACACGGCCCTCTT |
| VB_pr166_HsXBP1_L | AGCTTTTACGAGAGAAAACTCAT |
| VB_pr220_HsACT_RT_L | TTCTACAATGAGCTGCGTGTG |
| VB_pr221_HsACT_RT_R | AGGGACATACCCCTCGTAGAT |
| VB_pr222_HsXBP1s_RT_R | CCTGCACCTGCTGCG |
| VB_pr245_HsERdj4_L | GTCGGAGGGTGCAGGATATTAG |
| VB_pr246_HsERdj4_R | TCAGGGTGGTACTTCATGGC |
| VB_pr255_HsCHOP_L | CCTCCTGGAAATGAAGAGGAAGA |
| VB_pr256_HsCHOP_R | TCCTGGTTCTCCCTTGGTCT |
| VB_pr257_HsXBP1_L | TAAGACAGCGCTTGGGGATG |
| VB_pr258_HsXBP1_R | TGTTCTGGAGGGGTGACAAC |
| VB_pr261_HsXBP1_L | ATGAGTGAGCTGGAACAGCAA |
| VB_pr262_HsXBP1_R | GGCCTCACTTCATTCCCCTTG |

Table S1. DNA Oligo Sequences Used.

| Gene ID | Category | Description |
|---|---|---|
| MSL2 | chromatin | part of MSL complex, which regulates histone ubiquitylation. May also help respond to DNA damage.[1–3] |
| DLG2 | Other | seems to be a scaffold protein in signal transduction complexes in certain neurons[4,5] |
| RBL1 | chromatin | regulates chromatin modification. Related to retinoblastoma 1 (RB1), and controls cell cycle progression to some extent.[6–8] |
| TCF20 | transcription | transcription factor that seems to be important in neural development[9,10] |
| FBXL20 | Other | F-box protein. One of the substrate recognizing proteins in E3 ubiquitin ligases[11,12] |
| GATAD2B | chromatin | critical in the NURD histone deacetylation complex which represses genes.[13] |
| KANSL1 | chromatin | unstructured scaffold protein for the nonspecific lethal NSL complex that regulates histone modification, necessary for recruitment of WDR5. necessary for proper maintenance of epigenetic cell identity. Also important for mitochondrial function [14–16] |
| EIF4G3 | Other | part of eIF4F cap-binding complex of ribosome[17] |

| | | |
|---|---|---|
| NPIPB12 | Other | unknown function. |
| MED13 | transcription | component of the mediator complex, which is heavily involved in RNA PolII transcription[18,19] |
| SLC16A4 | Other | transmembrane solute transporter in the [20]monocarboxylate transporter family |
| LINC01297 | Other | uncharacterized lncRNA |
| DENND5B | lipid metabolism | Rab12 GEF[21,22] |
| MBTD1 | chromatin | Polycomb group protein. Binds to select histone modifications to regulate DNA repair pathways and transcriptional regulation [23,24] |
| DIP2B | Other | Possibly involved in DNA methylation, but there's not enough data to be confident.[25] |
| ZNF850 | Other | Probably a DNA binding protein, possibly involved in CTG repeat expansion[26] |
| LINC-PINT | Other | tumor/proliferation suppressor[27] |
| TLK2 | chromatin | Chromatin repair/maintenance[28] |
| NCOA6 | transcription | coregulator of transcriptional regulators and histone modifiers[29] |
| LOC100190986 | Other | uncharacterized transcript |
| CAMK1D | Other | potential regulator of Ca levels or signaling[30,31] |
| ASAP1 | Actin | GTPase involved in active cytoskeleton remodeling, especially at motile edges/podosomes[32] |
| TYW1B | Other | creates wybutosine residues on tRNa.[33] |
| ZFHX3 | transcription | repressive TF. cooperates with Smad2/3 to repress AFP, downstream of TGFbeta. Upregulated under hypoxia[34,35] |
| ARHGAP11B | mitochondrial function | inhibits adenine nucleotide translocase[36] |
| ARID1B | transcription | changes the gene targets of the chromatin remodelers SWI/SNF/BAF and inhibits Wnt/β-catenin. Important for proper cell differentiation and development. [37,38] |
| LINC01876 | Other | uncharacterized lncRNA |
| LOC442028 | Other | uncharacterized |
| CUX1 | transcription | TF involved with proliferation and possibly DNA damage repair[39] |
| MIR34AHG | transcription | lncRNA associated with ER stress. Possibly is a precursor to miR-34a, which seems to have broad regulatory effects, including p53 repression.[40,41] |
| TMEM120B | lipid metabolism | encourages adipogenesis[42] |
| RASA4 | Other | Under high intracellular Ca, Deactivates Ras, represses cancer (and other cell activities)[43,44] |
| DGAT2 | lipid metabolism | catalyzes DAG reaction with acyl-CoAs to make trigylcerides. Seems to have overlapping role with DGAT1 for TG synthesis, but they are unrelated proteins with different affinities for substrates and cofactors, and different regulation and activities[45,46] |

| | | |
|---|---|---|
| PVT1 | transcription | seems to upregulate c-Myc and increase proliferation[47] |
| ITGB5 | transcription | promotes metastisis and cell migration, maybe through Wnt/B-catenin pathway, smad, or TGFB[48,49] |
| FRMD5 | Actin | Interacts with integrins[50] |
| AGO2 | transcription | key part of the RISC silencing complex. Regulates chromatin modification through siRNA[51] |
| SVIL | Actin | large actin-binding protein that plays a role in motility and cytokinesis.[52,53] |
| ZNF782 | Other | uncharacterized zinc finger protein |
| DOCK5 | Actin | GEF of Rac1, a Rho GTPase. regulator of cytokinesis and actin dynamics.[54,55] |
| KRT17 | Other | keratin protein |
| MGAT4B | glycosylation | glycan branching[56,57] |
| TMEM179B | mitochondrial function | ROS protection in/near mitochondia.[58] |
| PCBP1-AS1 | transcription | seems to promote proliferation in cancers[59] |
| ANKDD1A | Other | uncharacterized ankyrin repeat protein. [60] |
| EPB41L4A-AS1 | mitochondrial function | regulating glycolysis and glutaminolysis[61] |
| BCAM | Other | regulates integrin binding to laminin, thus altering cell adhesion[62,63] |
| TMEM165 | ER-Golgi function | possible Ca/Mn and H+ antiporter that is needed for production of certain glycans, pulls Ca or Mn into ER[64–66] |
| FOXN3 | transcription | seems to repress Myc and glycolysis. Seems to repress smad2-4 signaling. Seems to repress proliferation[67,68] |
| SIPA1L3 | Actin | plays some role in actin cytoskeleton regulation[69] |
| ND1 | mitochondrial function | critical component of mitochondia complex I, a component of the ETC. Can be downregulated in tumors to promote glycolysis over oxidative respiration[70,71] |
| MLLT10 | transcription | activating TF that is related to development and differentiation and can directly induce colorectal cancer by promoting proliferation and invasion[72] |
| ASXL1 | chromatin | Coordinates deubiquitylation of histones involved with polycomb silencing[73,74] |
| HMGA2 | transcription | transcription factor that is disregulated in some cancers[75,76] |
| PIGQ | lipid metabolism | essential for GPI synthesis[77] |
| TIMP2 | Other | inhibits matrix metalloproteases. Seems to be stress induced[78,79] |
| EEF1A2 | Other | Part of the ribosome, helps with delivery of aminoacyl tRNAs to the ribosome[80] |
| GCLM | Other | Important for glutathione synthesis, which is one of the main ROS scavengers[81] |

| | | |
|---|---|---|
| MRPL23 | mitochondrial function | mitochondrial ribosome protein[82] |
| ZYX | Actin | zinc-binding phosphoprotein that functions with focal adhesions and actin cytoskeleton. Important for actin organization and cell motility[83,84] |
| RALB | ER-Golgi function | GTPase that seems to signal downstream of Ras. Seems to be actively involved with endomembrane autophagy and cell invasion/motility. Also improves DSB repair after irradiation[85,86] |
| VANGL1 | Actin | scaffolding protein related to cell polarity and organization in development[87] |
| WAPL | chromatin | regulates cohesin binding, which is critical for both chromatin organization and chromosome segregation[88,89] |
| RFTN1 | Other | lipid raft protein important in B cell antigen receptor function.[90,91] |
| LRP10 | ER-Golgi function | ER-golgi localized transmembrane protein, potentially involved in trafficking of lipoproteins. Also potentially involved in lewy body diseases[92,93] |
| DCBLD2 | Other | an orphan receptor with several proposed affiliations with cancer proliferation.[94] |
| SLAIN2 | Actin | important for microtubule formation and organization, especially during interphase[95] |
| LAMP1 | Other | One isoform of the most critical membrane protein in lysosomes.[96] |
| CAVIN1 | Actin | Major component of caveolae structure[97] |
| TYRO3 | Other | A receptor tyrosine kinase with several proposed functions[98] |
| AGPS | lipid metabolism | converts acyl-glycerone-3-phosphate into alkyl-glycerone-3-phosphate. Important for ether lipid synthesis[99] |
| FAM220A | transcription | regulates STAT3, which is a transcription factor involved in several processes.[100] |
| POLR3D | transcription | part of RNA PolIII[101] |
| CCDC6 | Other | seems to be a scaffolding protein that helps with genotoxic response[102,103] |
| IBTK | Other | inhibits BTK kinase, which is critical in activation of B cells downstream of B-cell antigen receptor via calcium release from the ER. Also activates NFkB[104] |
| ACTR2 | Actin | part of ARP2/3 complex, which is one of the actin nucleators that polymerize actin at the end of filaments. Partly responsible for cell motility[105] |
| FNDC3B | lipid metabolism | Important for adipocyte differentiation and other developmental processes. Also associated with UPR[106,107] |
| LTN1 | Other | ubiquitylates ribosomes stalled in non-stop translation on the polyA tail[108,109] |
| DNAJC3 | ER-Golgi function | co-chaperone for BiP[110] |
| HMOX1 | Other | heme oxygenase that is related to ferroptosis[111,112] |

| | | |
|---|---|---|
| OSBPL2 | lipid metabolism | lipid binding protein that likely regulates lipid transport and memrane composition, and may regulate lipid droplets in conjunction with the ER[113] |
| PGM3 | ER-Golgi function | enzyme necessary for creating UDP-GlcNAc, which is necessary for protein glycosylation[114] |
| ARMCX3 | mitochondrial function | plays some regulatory role regarding mitochondria regulation and trafficking, especially in neuronal development[115,116] |
| ALG2 | ER-Golgi function | mannosyltransferase important for glycosylation[117] |
| SRPRB | ER-Golgi function | component of the signal recognition particle receptor[118] |
| SLC33A1 | ER-Golgi function | membrane transporter of acetyl-CoA into the ER. Necessary for ganglioside acetylation[119,120] |
| FKBP14 | ER-Golgi function | Proline isomerase that helps folding in ER, especially for collagens[121,122] |
| GORASP2 | ER-Golgi function | Maintains golgi structure[123,124] |
| HERPUD1 | ER-Golgi function | ER membrane protein that is critical for ERAD[125,126] |
| FICD | ER-Golgi function | regulates BiP activity by AMPylating or deAMPylating it, depending on biological context.[127,128] |
| TRIM32 | Other | multifunctional protein with several proposed rolls[129] |
| DNAJB9 | ER-Golgi function | primarily ER chaperone that seems to either escort unfolded proteins to ERAD machinery or help them fold with BiP. Also gets localized to the nucleus in response to certain stresses.[130,131] |
| KLF10 | transcription | a repressor of TGF-β signaling[132] |
| SEC24D | ER-Golgi function | One paralog of SEC24, which is a critical component of COPII vessicles.[133] |
| GFPT1 | glycosylation | first and rate-limiting part of the hexosamine biosynthesis pathway[134,135] |
| TMED7 | ER-Golgi function | critical for ER-golgi translocation[136] |
| CANX | ER-Golgi function | ER chaperone[137,138] |
| DNAJC10 | ER-Golgi function | PDI in the ER[139] |
| SSR1 | ER-Golgi function | signal sequence receptor, helps with translocation into the ER during translation[140] |
| IQCG | Actin | seems to interact with calmodulin, maybe moderating calcium levels. Or it is a microtuble nucleator/regulator. Plays a role in sperm motility[141] |
| TMED10 | ER-Golgi function | critical for ER-golgi translocation. Seems to specifically bind GPI-anchored proteins. Co-regulated with TMED2 [136] |

| | | |
|---|---|---|
| CYB561 | mitochondrial function | membrane enzyme that does ascorbate redox, potentially involved in mitochondrial function[142] |
| TMED7-TICAM2 | ER-Golgi function | A readthrough transcript of TMED7, which is critical for ER-golgi translocation[136,143] |
| SEC61A1 | ER-Golgi function | part of the sec61 translocon that helps insert proteins into the ER membrane[144] |
| SSR3 | ER-Golgi function | signal sequence receptor, helps with translocation into the ER during translation[140] |
| SND1 | transcription | part of RISC complex. Also seems to alter alt splicing. Seems to regulate lipid metabolism genes.[145,146] |
| LMAN1 | ER-Golgi function | membrane-bound chaperone that is critical for making certain proteins like coagulation factors F-V and -VIII. Helps transport them to the Golgi[147,148] |
| PDIA3 | ER-Golgi function | PDI in the ER[149] |
| PLPP5 | lipid metabolism | enzyme that is involved with phospholipid metabolism[150] |
| ST6GALNAC4 | glycosylation | transfers sialic acid to glycan chains, preferrentially to glycoproteins[151,152] |
| FBXO16 | Other | F-box protein. One of the substrate recognizing proteins in E3 ubiquitin ligases. May repress beta catenin and inflammation[153,154] |
| OSTC | glycosylation | a subunit of the oligosaccharyltransferase complex, which acts near sec61 to glycosylate peptides. Upregulated under heat stress, seems to be specific to one version of the core of the OST complex[155,156] |
| SERP1 | ER-Golgi function | membrane protein associated with translation and translocation into the ER. Might help protect other membrane proteins from ERAD during ER stress[157,158] |
| CDK2AP2 | chromatin | inhibits cell cycle progression, also involved with the NuRD histone modification complex[159,160] |
| NANS | glycosylation | catalyzes a step in the pathway for sialic acid[161] |
| SEC61B | ER-Golgi function | part of the sec61 translocon that helps insert proteins into the ER membrane[144] |
| MYDGF | Other | seems to be a proliferative signal[162] |
| PPIB | ER-Golgi function | Part of a complex that functions as a PDI and to hydroxylate prolyl residues in preprocollagen. Works after P4HB[163] |
| SEC11C | ER-Golgi function | one of the paralogs of the catalytic subunit of SPC complex, which cleaves signal sequences in the ER[164] |
| TMED9 | ER-Golgi function | regulates COP vesicles to affect protein transportation between ER and Golgi[136] |
| NUCB2 | Other | multi-component peptide that is cleaved into 3 parts that seem to have different functions. Nesfatin-1 seems to be the most important and is used as a hunger-related hormone[165,166] |

| | | |
|---|---|---|
| SSR2 | ER-Golgi function | signal sequence receptor, helps with translocation into the ER during translation[140] |
| GMPPA | glycosylation | part of the complex that produces GDP-Mannose, which is necessary for mannose glycosylation. Seems to have a regulatory role while GMPPB is catalytic[167] |
| P4HB | ER-Golgi function | Part of a complex that functions as a PDI and to hydroxylate prolyl residues in preprocollagen. Works before PPIB[168] |
| HM13 | ER-Golgi function | one of the peptidases in ERAD that cleaves proteins from signal peptides during/after translocation. Has specific substrates[169] |
| ZSCAN31 | transcription | TF. Not very well studied, but seems to get regulated in many cancers[170] |
| TXNDC15 | ER-Golgi function | ER-resident thioredoxin related to the PDI family[171] |
| SMIM14 | Other | uncharacterized ER membrane protein[172] |

Table S2. Genes Found to be Regulated By Opto-IRE1.

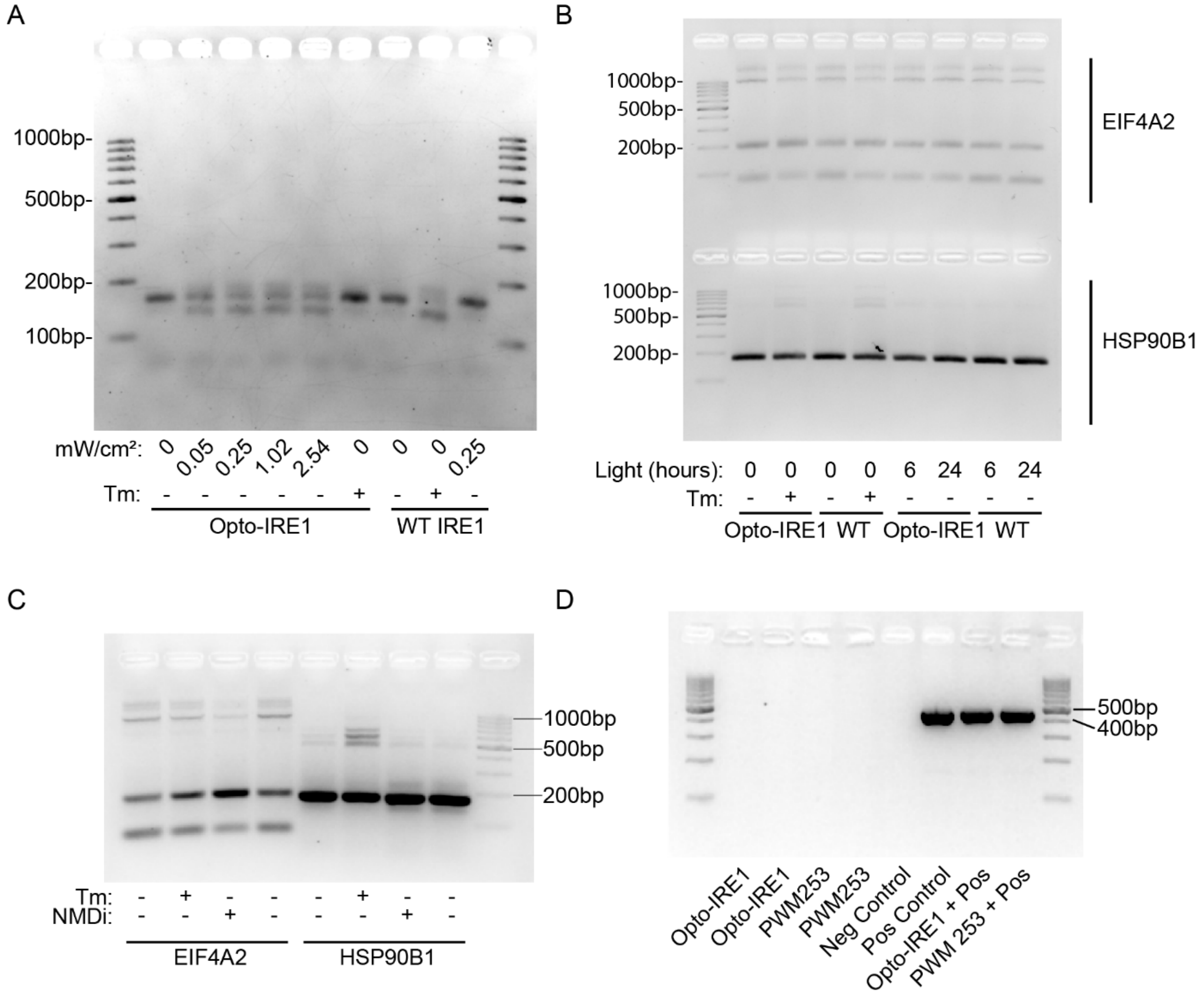


Figure S1. Uncropped Gels. (A) XBP1 splicing gel for Figure 1C. RNA was purified from Opto-IRE1 and WT IRE1 cells, reverse transcribed, and amplified with primers across the unconventional intron of XBP1 then resolved on a 3% agarose gel. (B) Gel of splicing assay for Figure 5, panels D and E. RNA was collected and reverse transcribed then amplified with primers targeting the alternatively spliced regions of the transcripts for *EIF4A2* (top half of gel) or *HSP90B1* (bottom half of gel). (C) Gel of splicing assay for Figure 6B. RNA was collected from WT cells and reverse transcribed then amplified with primers targeting the alternatively spliced regions of the transcripts for *EIF4A2* or *HSP90B1*. (D) Mycoplasma testing of Opto-IRE1 and WT cells. Cell lines were tested for mycoplasma contamination using Universal Mycoplasma Detection Kit (ATCC 301012K). The presence of a band at 434-468bp indicates the presence of mycoplasma DNA.

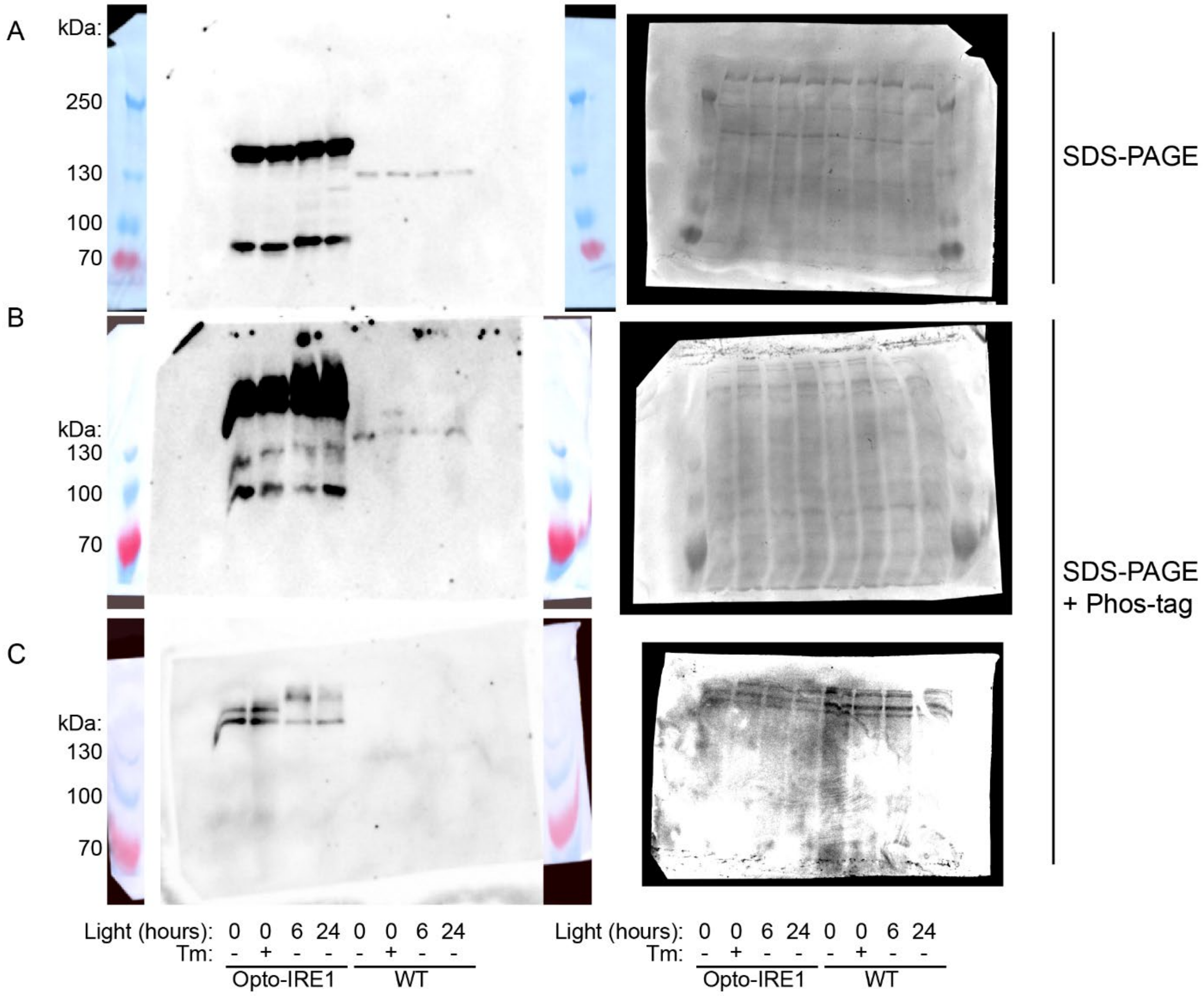


Fig S2. Uncropped Immunoblots for Figure 1B. Total protein was collected from Opto-IRE1 and WT IRE1 cells, separated by SDS-PAGE (A) or Phos-tag SDS-PAGE (B, C), and probed with anti-IRE1 antibody. 12µg of total protein was loaded per lane in panels A and B, while 10µg of total protein was loaded per lane in panel C to reduce the signal of the Opto-IRE1 bands. The WT lanes of panel B and the Opto-IRE1 lanes of panel C were used for Fig. 1B. Ponceau staining of each blot is shown on the right as a loading control.

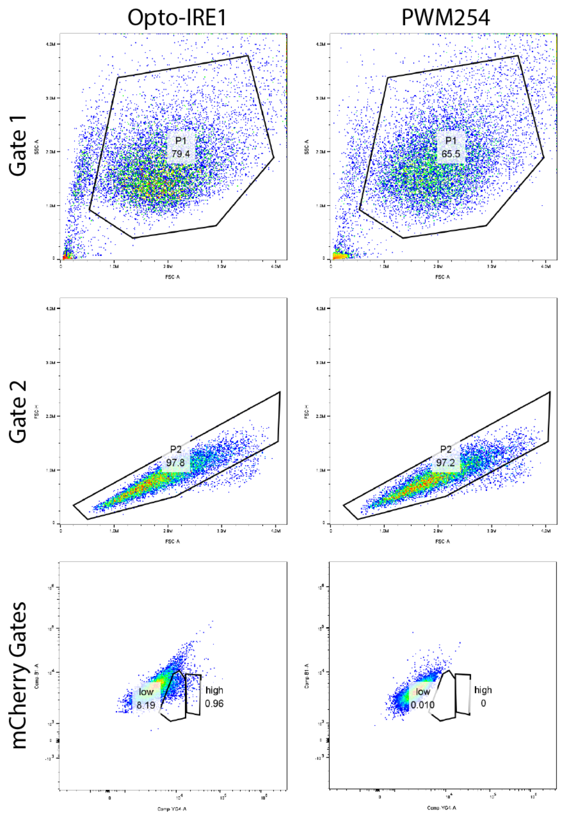


Fig S3. FACS Sorting of Opto-IRE1 and PWM254 cells. Droplets were gated twice to target single cells and then gated based on mCherry signal into low- and high-expressing populations.

A

```
Q: HSP90B1 intron-retaining sequence
S: GRCh38.p14 Chromosome 12

Q 103943266  AAAGAATTTGAGCCTCTGCTGAATTGGATGAAAGATAAAGCCCTTAAGGACAAGGTACTG  103943325
             |||||||||||||||||||||| ||||||||| |||||||||||||||||||||||||||
S 4          AAAGAATTTGAGCCTCTGCTGA-TTGGATGAA-GATAAAGCCCTTAAGGACAAGGTACTG  61

Q 103943326  TGGAAATTACAAATTGTGGAAATATTAGTATCAGCATTTAAGAGAAAGTTATTTTGTGAA  103943385
             ||||||||||||||||||||||||||||||||||||||||||||||||||||||||||||
S 62         TGGAAATTACAAATTGTGGAAATATTAGTATCAGCATTTAAGAGAAAGTTATTTTGTGAA  121

Q 103943386  CAAATTAAGCTGCAGCTGGTTACTTTGTAACCATTAGAATGGTAAAAATTTAATTAATGT  103943445
             ||||||||||||||||||||||||||||||||||||||||||||||||||||||||||||
S 122        CAAATTAAGCTGCAGCTGGTTACTTTGTAACCATTAGAATGGTAAAAATTTAATTAATGT  181

Q 103943446  AATTAAATTATTGGGAGAAAGCTTAAAACTTTCGACAATACTGCTTTGTTAATAACTTGT  103943505
             ||||||||||||||||||||||||||||||||||||||||||||||||||||||||||||
S 182        AATTAAATTATTGGGAGAAAGCTTAAAACTTTCGACAATACTGCTTTGTTAATAACTTGT  241

Q 103943506  TACAAATTAAATTTTATGTTTTTAAAAGGTGGTATTTAAACTTCTGACTAGAAAATTCAG  103943565
             ||||||||||||||||||||||||||||||||||||||||||||||||||||||||||||
S 242        TACAAATTAAATTTTATGTTTTTAAAAGGTGGTATTTAAACTTCTGACTAGAAAATTCAG  301

Q 103943566  ATTATCAAGTAAGTGCCCCTACAAATTCTCCTAAACCTTAAGAAAAGCTATTTTTATGAC  103943625
             ||||||||||||||||||||||||||||||||||||||||||||||||||||||||||||
S 302        ATTATCAAGTAAGTGCCCCTACAAATTCTCCTAAACCTTAAGAAAAGCTATTTTTATGAC  361

Q 103943626  CTGCTTCTGTGTTTATGATCTTAAGTGATAAAGTCTTAGACAGTTGAAAGACAATTGCTC  103943685
             ||||||||||||||||||||||||||||||||||||||||||||||||||||||||||||
S 362        CTGCTTCTGTGTTTATGATCTTAAGTGATAAAGTCTTAGACAGTTGAAAGACAATTGCTC  421

Q 103943686  AATGACCTTACCTGTTGATATTAATTTATATGACTTGATTTCTTTCCCTAAGATTGAAAA  103943745
             ||||||||||||||||||||||||||||||||||||||||||||||||||||||||||||
S 422        AATGACCTTACCTGTTGATATTAATTTATATGACTTGATTTCTTTCCCTAAGATTGAAAA  481

Q 103943746  GGCTGTGGTGTCTCAGCGCCTGACAGAATCTCCGTGTGCTTTGGTGGCCAGCCAGTACGG  103943805
             ||||||||||||||||||||||||||||||||||||||||||||||||||||||||||||
S 482        GGCTGTGGTGTCTCAGCGCCTGACAGAATCTCCGTGTGCTTTGGTGGCCAGCCAGTACGG  541

Q 103943806  ATGGTCTGGCAACATGGAGAGAATCATGAAA  103943836
             ||||||||||||||||||||| |||||| ||
S 542        ATGGTCTGGCAACATGGAGAG-ATCATGGAA  571
```

B

```
Q: EIF4A2 intron-retaining sequence
S: GRCh38.p14 Chomosome 3

Q 8          ACCCATCGTGAA--CTATATTCACAG  31
             ||| ||||||||  ||||||||||||
S 186787857  ACCAATCGTGAAAACTATATTCACAG  1867878829

Q 32         GAGTCGATAGCAGCAGTTGGTGACGAGATGGCACTCAGAAACGGCGTTGACGTAAT  87
             ||||||||||||||||||||||||||||||||||||||||||||||||||||||||
S  186788310 GAGTCGATAGCAGCAGTTGGTGACGAGATGGCACTCAGAAACGGCGTTGACGTAAT  186788365

Q 88         TTAGGACGTGGAATCATAAGCGAAACAGCACACTGTTTGAATAAAGAGCGAGTCG  142
             |||||||||||||||||||||||||||||||||||||||||||||||||||||||
S 186788366  TTAGGACGTGGAATCATAAGCGAAACAGCACACTGTTTGAATAAAGAGCGAGTCG  186788420
```

Fig S4. Sequencing Results of the Intron-retaining PCR Products for *EIF4A2* and *HSP90B1*. For EIF4A2, the ~200bp band was purified, sequenced, and aligned to the human genome. For *HSP90B1*, the ~600bp band was purified, sequenced, and aligned to the human genome using BLAST[173,174].

(114) Greig, K. T.; Antonchuk, J.; Metcalf, D.; Morgan, P. O.; Krebs, D. L.; Zhang, J.-G.; Hacking, D. F.; Bode, L.; Robb, L.; Kranz, C.; de Graaf, C.; Bahlo, M.; Nicola, N. A.; Nutt, S. L.; Freeze, H. H.; Alexander, W. S.; Hilton, D. J.; Kile, B. T. Agm1/Pgm3-Mediated Sugar Nucleotide Synthesis Is

Essential for Hematopoiesis and Development. *Mol. Cell. Biol.* **2007**, *27* (16), 5849–5859. https://doi.org/10.1128/MCB.00802-07.
(115) López-Doménech, G.; Serrat, R.; Mirra, S.; D'Aniello, S.; Somorjai, I.; Abad, A.; Vitureira, N.; García-Arumí, E.; Alonso, M. T.; Rodriguez-Prados, M.; Burgaya, F.; Andreu, A. L.; García-Sancho, J.; Trullas, R.; Garcia-Fernàndez, J.; Soriano, E. The Eutherian Armcx Genes Regulate Mitochondrial Trafficking in Neurons and Interact with Miro and Trak2. *Nat. Commun.* **2012**, *3* (1), 814. https://doi.org/10.1038/ncomms1829.
(116) Izquierdo-Villalba, I.; Mirra, S.; Manso, Y.; Parcerisas, A.; Rubio, J.; Del Valle, J.; Gil-Bea, F. J.; Ulloa, F.; Herrero-Lorenzo, M.; Verdaguer, E.; Benincá, C.; Castro-Torres, R. D.; Rebollo, E.; Marfany, G.; Auladell, C.; Navarro, X.; Enríquez, J. A.; López de Munain, A.; Soriano, E.; Aragay, A. M. A Mammalian-Specific Alex3/Gαq Protein Complex Regulates Mitochondrial Trafficking, Dendritic Complexity, and Neuronal Survival. *Sci. Signal.* **2024**, *17* (822), eabq1007. https://doi.org/10.1126/scisignal.abq1007.
(117) O'Reilly, M. K.; Zhang, G.; Imperiali, B. In Vitro Evidence for the Dual Function of Alg2 and Alg11: Essential Mannosyltransferases in N-Linked Glycoprotein Biosynthesis. *Biochemistry* **2006**, *45* (31), 9593–9603. https://doi.org/10.1021/bi060878o.
(118) Schwartz, T.; Blobel, G. Structural Basis for the Function of the β Subunit of the Eukaryotic Signal Recognition Particle Receptor. *Cell* **2003**, *112* (6), 793–803. https://doi.org/10.1016/S0092-8674(03)00161-2.
(119) Kanamori, A.; Nakayama, J.; Fukuda, M. N.; Stallcup, W. B.; Sasaki, K.; Fukuda, M.; Hirabayashi, Y. Expression Cloning and Characterization of a cDNA Encoding a Novel Membrane Protein Required for the Formation of O-Acetylated Ganglioside: A Putative Acetyl-CoA Transporter. *Proc. Natl. Acad. Sci. U. S. A.* **1997**, *94* (7), 2897–2902. https://doi.org/10.1073/pnas.94.7.2897.
(120) Jonas, M. C.; Pehar, M.; Puglielli, L. AT-1 Is the ER Membrane Acetyl-CoA Transporter and Is Essential for Cell Viability. *J. Cell Sci.* **2010**, *123* (Pt 19), 3378–3388. https://doi.org/10.1242/jcs.068841.
(121) Ishikawa, Y.; Bächinger, H. P. A Substrate Preference for the Rough Endoplasmic Reticulum Resident Protein FKBP22 during Collagen Biosynthesis. *J. Biol. Chem.* **2014**, *289* (26), 18189–18201. https://doi.org/10.1074/jbc.M114.561944.
(122) Boudko, S. P.; Ishikawa, Y.; Nix, J.; Chapman, M. S.; Bächinger, H. P. Structure of Human Peptidyl-Prolyl Cis–Trans Isomerase FKBP22 Containing Two EF-Hand Motifs. *Protein Sci.* **2014**, *23* (1), 67–75. https://doi.org/10.1002/pro.2391.
(123) Zhang, Y.; Seemann, J. Rapid Degradation of GRASP55 and GRASP65 Reveals Their Immediate Impact on the Golgi Structure. *J. Cell Biol.* **2020**, *220* (1), e202007052. https://doi.org/10.1083/jcb.202007052.
(124) Grond, R.; Veenendaal, T.; Duran, J. M.; Raote, I.; van Es, J. H.; Corstjens, S.; Delfgou, L.; El Haddouti, B.; Malhotra, V.; Rabouille, C. The Function of GORASPs in Golgi Apparatus Organization in Vivo. *J. Cell Biol.* **2020**, *219* (9), e202004191. https://doi.org/10.1083/jcb.202004191.
(125) Huang, C.-H.; Chu, Y.-R.; Ye, Y.; Chen, X. Role of HERP and a HERP-Related Protein in HRD1-Dependent Protein Degradation at the Endoplasmic Reticulum*. *J. Biol. Chem.* **2014**, *289* (7), 4444–4454. https://doi.org/10.1074/jbc.M113.519561.
(126) Okuda-Shimizu, Y.; Hendershot, L. M. Characterization of an ERAD Pathway for Nonglycosylated BiP Substrates, Which Require Herp. *Mol. Cell* **2007**, *28* (4), 544–554. https://doi.org/10.1016/j.molcel.2007.09.012.
(127) Casey, A. K.; Stewart, N. M.; Zaidi, N.; Gray, H. F.; Cox, A.; Fields, H. A.; Orth, K. FicD Regulates Adaptation to the Unfolded Protein Response in the Murine Liver. *Biochimie* **2024**, *225*, 114–124. https://doi.org/10.1016/j.biochi.2024.05.012.

(128) Preissler, S.; Rato, C.; Perera, L. A.; Saudek, V.; Ron, D. FICD Acts Bifunctionally to AMPylate and De-AMPylate the Endoplasmic Reticulum Chaperone BiP. *Nat. Struct. Mol. Biol.* **2017**, *24* (1), 23–29. https://doi.org/10.1038/nsmb.3337.
(129) Bawa, S.; Piccirillo, R.; Geisbrecht, E. R. TRIM32: A Multifunctional Protein Involved in Muscle Homeostasis, Glucose Metabolism, and Tumorigenesis. *Biomolecules* **2021**, *11* (3), 408. https://doi.org/10.3390/biom11030408.
(130) Daverkausen-Fischer, L.; Pröls, F. The Function of the Co-Chaperone ERdj4 in Diverse (Patho-)Physiological Conditions. *Cell. Mol. Life Sci.* **2021**, *79* (1), 9. https://doi.org/10.1007/s00018-021-04082-4.
(131) Lai, C. W.; Otero, J. H.; Hendershot, L. M.; Snapp, E. ERdj4 Protein Is a Soluble Endoplasmic Reticulum (ER) DnaJ Family Protein That Interacts with ER-Associated Degradation Machinery *. *J. Biol. Chem.* **2012**, *287* (11), 7969–7978. https://doi.org/10.1074/jbc.M111.311290.
(132) Memon, A.; Lee, W. K. KLF10 as a Tumor Suppressor Gene and Its TGF-β Signaling. *Cancers* **2018**, *10* (6), 161. https://doi.org/10.3390/cancers10060161.
(133) Chatterjee, S.; Choi, A. J.; Frankel, G. A Systematic Review of Sec24 Cargo Interactome. *Traffic* **2021**, *22* (12), 412–424. https://doi.org/10.1111/tra.12817.
(134) Marshall, S.; Bacote, V.; Traxinger, R. R. Discovery of a Metabolic Pathway Mediating Glucose-Induced Desensitization of the Glucose Transport System. Role of Hexosamine Biosynthesis in the Induction of Insulin Resistance. *J. Biol. Chem.* **1991**, *266* (8), 4706–4712.
(135) Ruegenberg, S.; Horn, M.; Pichlo, C.; Allmeroth, K.; Baumann, U.; Denzel, M. S. Loss of GFAT-1 Feedback Regulation Activates the Hexosamine Pathway That Modulates Protein Homeostasis. *Nat. Commun.* **2020**, *11* (1), 687. https://doi.org/10.1038/s41467-020-14524-5.
(136) Zhou, L.; Li, H.; Yao, H.; Dai, X.; Gao, P.; Cheng, H. TMED Family Genes and Their Roles in Human Diseases. *Int. J. Med. Sci.* **2023**, *20* (13), 1732–1743. https://doi.org/10.7150/ijms.87272.
(137) Bergeron, J. J. M.; Brenner, M. B.; Thomas, D. Y.; Williams, D. B. Calnexin: A Membrane-Bound Chaperone of the Endoplasmic Reticulum. *Trends Biochem. Sci.* **1994**, *19* (3), 124–128. https://doi.org/10.1016/0968-0004(94)90205-4.
(138) Paskevicius, T.; Farraj, R. A.; Michalak, M.; Agellon, L. B. Calnexin, More Than Just a Molecular Chaperone. *Cells* **2023**, *12* (3), 403. https://doi.org/10.3390/cells12030403.
(139) Robinson, P. J.; Pringle, M. A.; Fleming, B.; Bulleid, N. J. Distinct Role of ERp57 and ERdj5 as a Disulfide Isomerase and Reductase during ER Protein Folding. *J. Cell Sci.* **2023**, *136* (2), jcs260656. https://doi.org/10.1242/jcs.260656.
(140) Russo, A. Understanding the Mammalian TRAP Complex Function(s). *Open Biol.* **2020**, *10* (5), 190244. https://doi.org/10.1098/rsob.190244.
(141) Li, R.-K.; Tan, J.-L.; Chen, L.-T.; Feng, J.-S.; Liang, W.-X.; Guo, X.-J.; Liu, P.; Chen, Z.; Sha, J.-H.; Wang, Y.-F.; Chen, S.-J. Iqcg Is Essential for Sperm Flagellum Formation in Mice. *PLOS ONE* **2014**, *9* (5), e98053. https://doi.org/10.1371/journal.pone.0098053.
(142) Asard, H.; Barbaro, R.; Trost, P.; Bérczi, A. Cytochromes B561: Ascorbate-Mediated Trans-Membrane Electron Transport. *Antioxid. Redox Signal.* **2013**, *19* (9), 1026–1035. https://doi.org/10.1089/ars.2012.5065.
(143) Doyle, S. L.; Husebye, H.; Connolly, D. J.; Espevik, T.; O’Neill, L. A. J.; McGettrick, A. F. The GOLD Domain-Containing Protein TMED7 Inhibits TLR4 Signalling from the Endosome upon LPS Stimulation. *Nat. Commun.* **2012**, *3* (1), 707. https://doi.org/10.1038/ncomms1706.
(144) Lang, S.; Pfeffer, S.; Lee, P.-H.; Cavalié, A.; Helms, V.; Förster, F.; Zimmermann, R. An Update on Sec61 Channel Functions, Mechanisms, and Related Diseases. *Front. Physiol.* **2017**, *8*. https://doi.org/10.3389/fphys.2017.00887.
(145) Cappellari, M.; Bielli, P.; Paronetto, M. P.; Ciccosanti, F.; Fimia, G. M.; Saarikettu, J.; Silvennoinen, O.; Sette, C. The Transcriptional Co-Activator SND1 Is a Novel Regulator of Alternative Splicing in

Prostate Cancer Cells. *Oncogene* **2014**, *33* (29), 3794–3802. https://doi.org/10.1038/onc.2013.360.
(146) Navarro-Imaz, H.; Ochoa, B.; García-Arcos, I.; Martínez, M. J.; Chico, Y.; Fresnedo, O.; Rueda, Y. Molecular and Cellular Insights into the Role of SND1 in Lipid Metabolism. *Biochim. Biophys. Acta BBA - Mol. Cell Biol. Lipids* **2020**, *1865* (5), 158589. https://doi.org/10.1016/j.bbalip.2019.158589.
(147) Cunningham, M. A.; Pipe, S. W.; Zhang, B.; Hauri, H.-P.; Ginsburg, D.; Kaufman, R. J. LMAN1 Is a Molecular Chaperone for the Secretion of Coagulation Factor VIII1. *J. Thromb. Haemost.* **2003**, *1* (11), 2360–2367. https://doi.org/10.1046/j.1538-7836.2003.00415.x.
(148) Zheng, C.; Liu, H.; Yuan, S.; Zhou, J.; Zhang, B. Molecular Basis of LMAN1 in Coordinating LMAN1-MCFD2 Cargo Receptor Formation and ER-to-Golgi Transport of FV/FVIII. *Blood* **2010**, *116* (25), 5698–5706. https://doi.org/10.1182/blood-2010-04-278325.
(149) Mahmood, F.; Xu, R.; Awan, M. U. N.; Song, Y.; Han, Q.; Xia, X.; Zhang, J. PDIA3: Structure, Functions and Its Potential Role in Viral Infections. *Biomed. Pharmacother.* **2021**, *143*, 112110. https://doi.org/10.1016/j.biopha.2021.112110.
(150) Takeuchi, M.; Harigai, M.; Momohara, S.; Ball, E.; Abe, J.; Furuichi, K.; Kamatani, N. Cloning and Characterization of DPPL1 and DPPL2, Representatives of a Novel Type of Mammalian Phosphatidate Phosphatase. *Gene* **2007**, *399* (2), 174–180. https://doi.org/10.1016/j.gene.2007.05.009.
(151) Man, D.; Jiang, Y.; Zhang, D.; Wu, J.; Ding, B.; Liu, H.; Xu, G.; Lu, J.; Ru, J.; Tong, R.; Zheng, S.; Chen, D.; Wu, J. ST6GALNAC4 Promotes Hepatocellular Carcinogenesis by Inducing Abnormal Glycosylation. *J. Transl. Med.* **2023**, *21* (1), 420. https://doi.org/10.1186/s12967-023-04191-7.
(152) Isaji, T.; Gu, J. Novel Regulatory Mechanisms of N-Glycan Sialylation: Implication of Integrin and Focal Adhesion Kinase in the Regulation. *Biochim. Biophys. Acta BBA - Gen. Subj.* **2024**, *1868* (6), 130617. https://doi.org/10.1016/j.bbagen.2024.130617.
(153) Paul, D.; Islam, S.; Manne, R. K.; Dinesh, U.; Malonia, S. K.; Maity, B.; Boppana, R.; Rapole, S.; Shetty, P. K.; Santra, M. K. F-Box Protein FBXO16 Functions as a Tumor Suppressor by Attenuating Nuclear β-Catenin Function. *J. Pathol.* **2019**, *248* (3), 266–279. https://doi.org/10.1002/path.5252.
(154) Sugimoto-Ishige, A.; Jodo, A.; Tanaka, T. Fbxo16 Mediates Degradation of NF-κB P65 Subunit and Inhibits Inflammatory Response in Dendritic Cells. *Front. Immunol.* **2025**, *16*. https://doi.org/10.3389/fimmu.2025.1524110.
(155) Chen, C.-M.; Tseng, C.-N.; Cho, J. J.; Lee, Y.-Z.; Kao, C.-L.; Cheng, Y.-B.; Hong, Y.-R.; Cho, C.-L. Heat Shock Induces Expression of OSTC/DC2, a Novel Subunit of Oligosaccharyltransferase, *in Vitro* and *in Vivo*. *Kaohsiung J. Med. Sci.* **2014**, *30* (5), 219–223. https://doi.org/10.1016/j.kjms.2014.01.003.
(156) Shrimal, S.; Cherepanova, N. A.; Gilmore, R. DC2 and KCP2 Mediate the Interaction between the Oligosaccharyltransferase and the ER Translocon. *J. Cell Biol.* **2017**, *216* (11), 3625–3638. https://doi.org/10.1083/jcb.201702159.
(157) Jung, S.; Hyun, J.; Nah, J.; Han, J.; Kim, S.-H.; Park, J.; Oh, Y.; Gwon, Y.; Moon, S.; Jo, D.-G.; Jung, Y.-K. SERP1 Is an Assembly Regulator of γ-Secretase in Metabolic Stress Conditions. *Sci. Signal.* **2020**, *13* (623), eaax8949. https://doi.org/10.1126/scisignal.aax8949.
(158) Yamaguchi, A.; Hori, O.; Stern, D. M.; Hartmann, E.; Ogawa, S.; Tohyama, M. Stress-Associated Endoplasmic Reticulum Protein 1 (Serp1)/Ribosome-Associated Membrane Protein 4 (Ramp4) Stabilizes Membrane Proteins during Stress and Facilitates Subsequent Glycosylation. *J. Cell Biol.* **1999**, *147* (6), 1195–1204. https://doi.org/10.1083/jcb.147.6.1195.
(159) Spruijt, C. G.; Gräwe, C.; Kleinendorst, S. C.; Baltissen, M. P. A.; Vermeulen, M. Cross-Linking Mass Spectrometry Reveals the Structural Topology of Peripheral NuRD Subunits Relative to the Core Complex. *FEBS J.* **2021**, *288* (10), 3231–3245. https://doi.org/10.1111/febs.15650.
